# The persimmon genome reveals clues to the evolution of a lineage-specific sex determination system in plants

**DOI:** 10.1101/628537

**Authors:** Takashi Akagi, Kenta Shirasawa, Hideki Nagasaki, Hideki Hirakawa, Ryutaro Tao, Luca Comai, Isabelle M. Henry

## Abstract

Most angiosperms bear hermaphroditic flowers, but a few species have evolved outcrossing strategies, such as dioecy, the presence of separate male and female individuals. We previously investigated the mechanisms underlying dioecy in diploid persimmon (*D. lotus*) and found that male flowers are specified by repression of the autosomal gene *MeGI* by its paralog, the Y-encoded pseudo-gene *OGI*. This mechanism is thought to be lineage-specific, but its evolutionary path remains unknown. Here, we developed a full draft of the diploid persimmon genome (*D. lotus*), which revealed a lineage-specific genome-wide paleoduplication event. Together with a subsequent persimmon-specific duplication(s), these events resulted in the presence of three paralogs, *MeGI*, *OGI* and newly identified *Sister of MeGI* (*SiMeGI*), from the single original gene. Evolutionary analysis suggested that *MeGI* underwent adaptive evolution after the paleoduplication event. Transformation of tobacco plants with *MeGI* and *SiMeGI* revealed that *MeGI* specifically acquired a new function as a repressor of male organ development, while *SiMeGI* presumably maintained the original function. Later, local duplication spawned *MeGI*’s regulator *OGI*, completing the path leading to dioecy. These findings exemplify how duplication events can provide flexible genetic material available to help respond to varying environments and provide interesting parallels for our understanding of the mechanisms underlying the transition into dieocy in plants.

**Author summary:** Plant sexuality has fascinated scientists for decades. Most plants can self-reproduce but not all. For example, a small subset of species have evolved a system called dioecy, with separate male and female individuals. Dioecy has evolved multiple times independently and, while we do not understand the molecular mechanisms underlying dioecy in many of these species yet, a picture is starting to emerge with recent progress in several dioecious species. Here, we focused on the evolutionary events leading to dioecy in persimmon. Our previous work had identified a pair of genes regulating sex in this species, called *OGI* and *MeGI*. We drafted the whole genome sequence of diploid persimmon to investigate their evolutionary history. We discovered a lineage-specific genome duplication event, and observed that *MeGI* underwent adaptive evolution after this duplication. Transgenic analyses validated that *MeGI* newly acquired a male-suppressor function, while the other copy of this gene, *SiMeGI*, did not. The regulator of *MeGI*, *OGI*, resulted from a second smaller-scale duplication event, finalizing the system. This study sheds light on the role of duplication as a mechanism that promote flexible genes functions, and how it can affect important biological functions, such as the establishment of a new sexual system.

## Introduction

Most species of flowering plants are hermaphrodite, but a small proportion have genetically determined separate sexes (Renner, 2014). The rarity of dioecy contrasts with its broad distribution across the flowering plant phylogenetic tree, suggesting multiple independent transitions into dioecy. Our study aimed to understand the molecular and evolutionary mechanisms underlying such changes. Advances in genomic analyses have allowed studies of plant sex chromosomes in a few dioecious plant species including papaya and *Silene* (Liu et al., 2004; Wang et al., 2012; Kazama et al., 2016), and a few genetic sex determining genes have recently been identified, including in the persimmon, kiwifruit, and asparagus (Akagi et al., 2014, 2018; Harkess et al., 2017). Consistent with theoretical models (Charlesworth and Charlesworth, 1978a, b), the results indicate that at least one gain-of-function mutation occurred in the evolution of dioecy, creating a dominant gynoecium or androecium suppressor. Data from these species is also consistent with gene duplication events as the first event leading to these gain-of-function mutations, because the redundancy provided by the presence of duplicate copies allows one copy to be neofunctionalized without loss of the original function (Flagel and Wendel, 2009). Unlike many animal taxa, flowering plants have experienced numerous whole-genome duplication events (WGD) (Van de Peer et al., 2017), which are thought to have provided opportunities for the appearance of new traits specific to each plant species. For example, functional differentiation between paralogs, which had been derived from whole-genome duplication (WGD), resulted in the establishment of ripening characteristics in tomato fruits (The Tomato Genome Consortium, 2012), and potentially enabled the adaptation to life underwater in seagrass (*Zostera marina*) (Olsen et al., 2016).

Within the large order Ericales, a heterogametic male (XY) sex determination system has evolved independently in at least two genera, *Diospyros* and *Actinidia* (Fraser et al., 2009; Akagi et al., 2014, 2018). *Diospyros* had evolved a Y-encoded pseudogene called *OGI*, that produces small-RNA, which in turn repress the autosomal feminization gene, *MeGI* (Akagi et al., 2014). *MeGI* belongs to the HD-Zip1 gene family conserved across angiosperms, but the specific function of *MeGI* to act for repression of male function, or feminization, has not been observed in *MeGI* orthologs from other plants so far (Komatsuda et al., 2007; Whipple et al., 2011; Sakuma et al., 2013). Indeed, although *Actinidia* and *Diospyros* are phylogenetically close to each other, the Y-encoded sex determination system in *Actinidia* does not involve the *MeGI* ortholog or another member of the HD-Zip1 family (Akagi et al., 2018). The existence of *MeGI*, *OGI*, and a third paralog called *Sister-of-MeGI* (*SiMeGI*), which was newly identified in this study, provide the opportunity to investigate both the scale and context of gene duplication events that triggered the appearance of a lineage-specific sex determination system in this species. To address this question, we sequenced the genome of Caucasian diploid persimmon, focusing on the lineage-specific duplication events. Evolutionary analyses on the duplicated pairs found a limited numbers of the genes which were potentially neofunctionalized via adaptive evolution after the duplication. Our results provide a potential path from the duplicated paralogs of a HD-Zip1 to dioecy, and shed light on how lineage-specific duplication events contribute to the evolution of a new sex determination system in a plant species.

## Results and Discussion

### Draft genome sequencing of Diospyros

Initially, we assembled a draft genome from ca 65X PacBio long read coverage of the expected haploid genome size (907Mb from nuclear weight (Tamura et al., 1998), 877.7Mb from kmer analysis) using Falcon (Supplemental Figure 1, Supplemental Table 1). This resulted in 3,073 primary contigs, and 5,901 “secondary” contigs, which are putative allelic contigs to the primary contigs. Next, we built three genetic maps, created from two segregating F1 populations (N = 314 and 119, see Materials and Methods and Supplemental Table 2). These maps were created from a total of 5,959 markers derived from GBS/ddRAD sequencing and allowed for the anchoring of the contigs into a genome draft comprised of 15 pseudomolecules (Figure 1, Supplemental Figure 1, Supplemental Tables 2 and 3).

**Figure 1.**
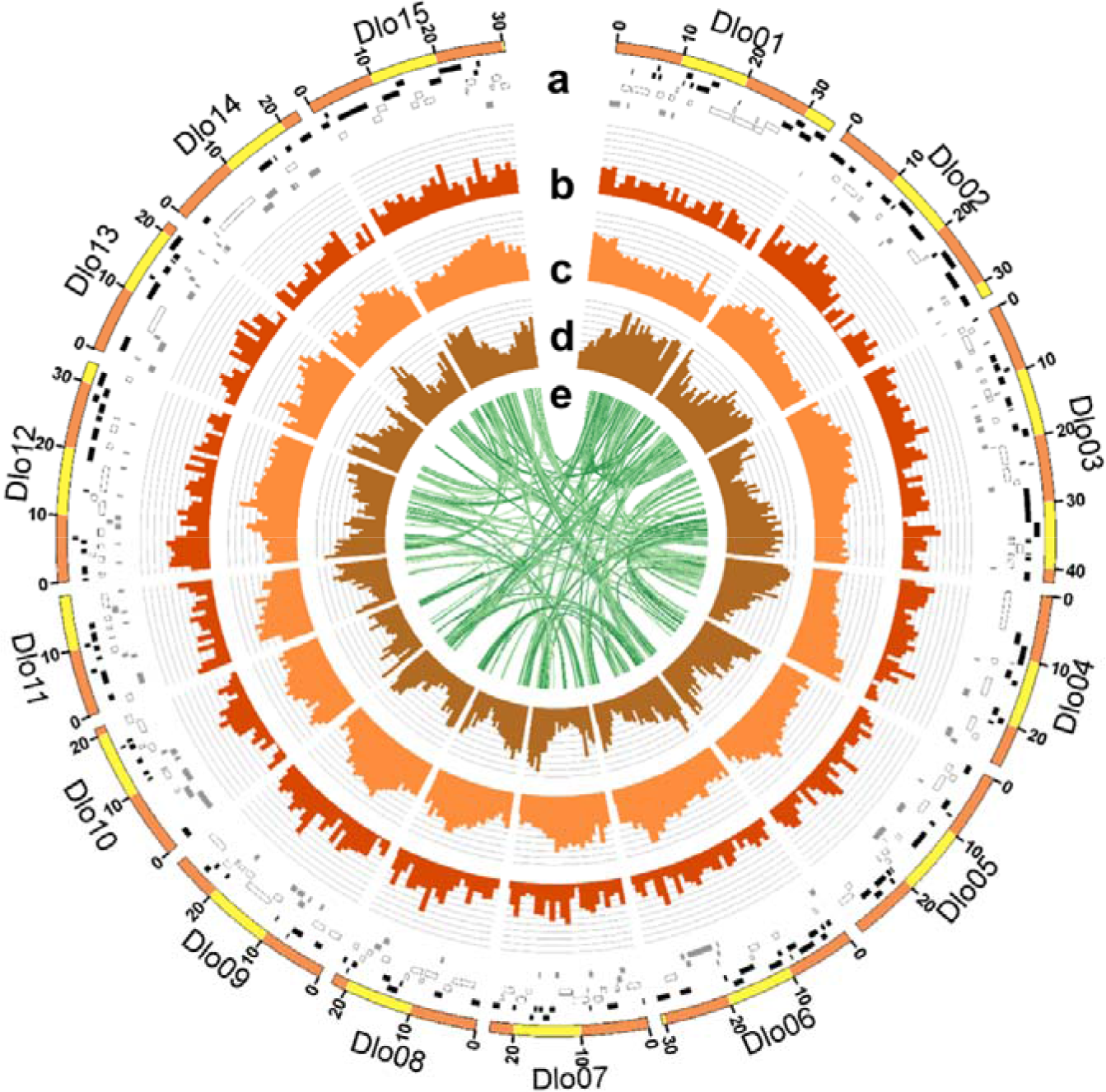
Characterization of the draft persimmon genome. **a**, Fifteen pseudomolecules with the genetically anchored contigs. Black, white and gray bars indicate the positions of the original contigs that were assembled in forward, reverse, or unknown direction respectively. **b**, Relative SNP density in the KK population. **c**, Relative density of repetitive sequences **d**, Relative gene density. **e**, Syntenic relationships within the persimmon genome.

To start characterizing this newly assembled genome, we documented sequence variation between female and male individuals of *D. lotus* and content and type of repeat sequences of the draft sequence compared to other sequences eudicots (Figure 1b and 1c, Supplemental Tables 3 and 4). Mapping of transcriptome data to this draft genome resulted in 40,532 predicted gene locations (Figure 1d, Supplemental Dataset 1). These numbers are similar to results from other asterid plant species, such as tomato (N = 34,879) (The Tomato Genome Consortium, 2012) or kiwifruit (N = 39,040) (Huang et al., 2013) (Supplemental Figures 2 and 3). Of these primary genes, we selected 12,058 which were determined to be either unique or low copy number within the genome (see Materials and Methods).

**Figure 2.**
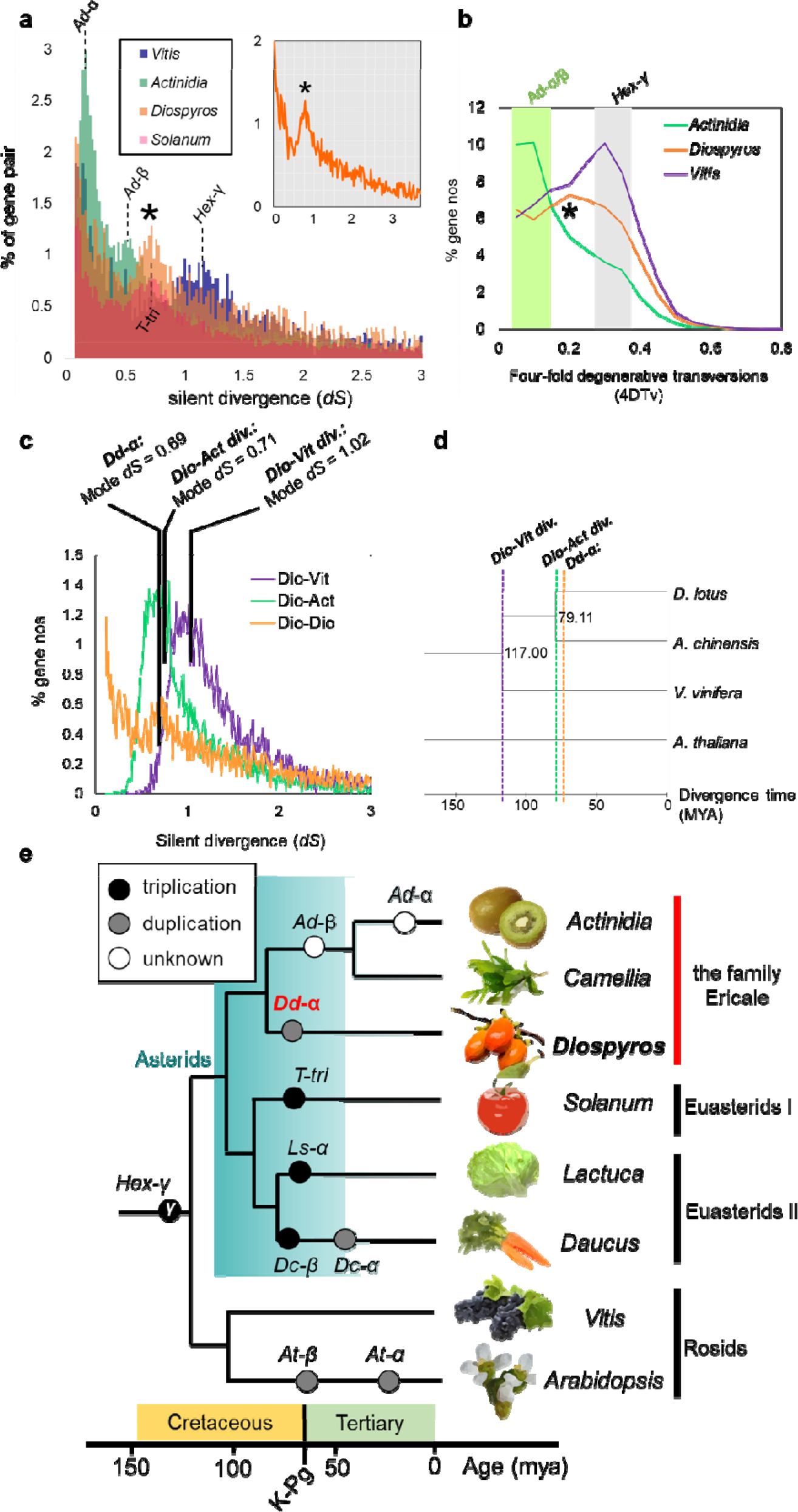
Characterization of lineage-specific whole-genome duplication events. **a**, Distribution of silent divergence rates between homologous gene pairs within the *Diospyros*, *Actinidia, Solanum*, and *Vitis* genomes. *Diospyros* shows a peak, indicated by an asterisk, at the same *dS* value as the *Solanum* triplication (T-tri), indicating the concurrent whole-genome duplication events. **b**, Comparison of the 4-fold degenerative transversion rates (4DTv) between the putative paralogous gene pairs, in the *Diospyros*, *Actinidia*, and *Vitis* genomes. Consistent with the distribution of *dS* values, a peak, which corresponds to *Dd-α*, was detected specifically in the persimmon genome, as indicated by an asterisk (*). In the *Actinidia* and *Vitis* genomes, peaks putatively corresponding the *Ad-α*/*β* and the hexaploidization-γ, were detected, as shown by the green and gray bands, respectively. **c**, Comparison of the *dS* values between the paralogous pairs in the *Diospyros* genome (orange), and the *dS* values between the orthologs in *Diospyros* and *Actinidia* (green), and in *Diospyros* and *Vitis* (purple). **d**, Estimated divergence time between *Diospyros*, *Actinidia*, and *Vitis*, with Arabidopsis as the outgroup. The concatenated sequences of 175 conserved genes across these species were used to determine divergence time, based on the previous estimated divergence of *Actinidia* and *Vitis* at 117MYA in the TIMETREE database (http://www.timetree.org). **e**, Summary of the lineage-specific WGD events in the asterids. The time scale is estimated from *dS* values and previous reports (Vanneste et al., 2014; Iorizzo et al., 2016; Reyes-Chin-Wo et al., 2017). K-Pg, Cretaceous-Paleogene boundary.

**Figure 3.**
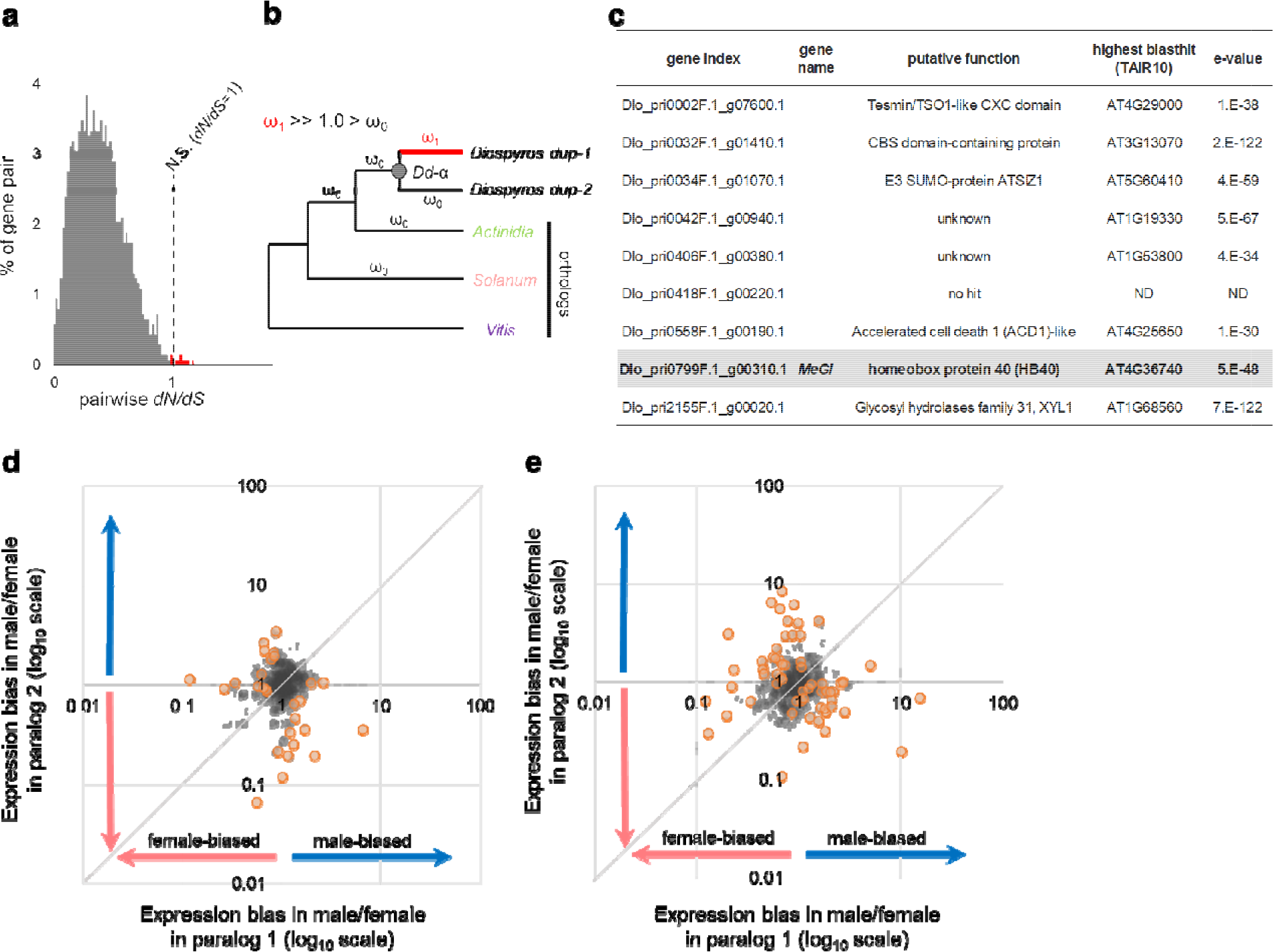
Fate of paleoduplicated genes in *Diospyros*. **a**, Distribution of the pairwise *dN/dS* values in the *Dd-α*-derived paralogous gene pairs, from the alignment of the full ORF sequences. Most gene pairs are under purifying selection (*dN/dS* < 1.0), while only approximately 0.3% of the gene pairs (shown in red) exhibited neutral selection (N.S.) or weak positive selection (*dN/dS* ~ 1.0). **b**, model for the detection of the genes that underwent significant site-branch specific positive selection (posterior probability > 0.99 in Bayes Empirical Bayes method) after *Dd-α*, using *Actinidia*, *Solanum*, and *Vitis* as outgroups. **c**, Functional annotation of the 9 genes that underwent significant site-branch specific positive selection after *Dd-α*. *MeGI* is highlighted in gray. **d-e**, Comparison of the expression patterns of paralog pairs derived from the *Dd-α* event, focusing on the sex differentiation stages. The ratio of expression levels in male versus female developing flowers (**d**) and mature flowers (**e**) were compared in the paralogs putatively derived from the *Dd-α* WGD event. The ratios were expressed in log_10_ scale. Approximately 10% of the gene pairs exhibited a statistically significant (*P* < 0.01, 2×2 Fisher’s exact test, orange circles) expression bias between the two paralogs (Supplemental Dataset 3), and 18.5% of the gene pairs (*N* = 242) showed >5-fold differences between the two paralogs.

### Identification of a whole-genome duplication event specific to the Diospyros genus

To investigate gene duplication patterns, we analyzed the distribution of silent divergence rate (*dS*) between homologous gene pairs. We compared the distribution of silent divergence rate of homologous gene pairs within the persimmon genome, with those within the kiwifruit (*Actinidia*), tomato (*Solanum*) and grape (*Vitis*) genomes. A subset of persimmon genes formed a clear peak of silent divergence rate (Figure 2a, *dS* = ca 0.5-0.9, mode *dS* = 0.69), suggesting that a whole-genome duplication (WGD) event, named *Dd-α*, occurred in this clade, and approximately simultaneously with the tomato genome triplication (The Tomato Genome Consortium, 2012) (Figure 2a). The genomic regions including the gene pairs in this peak exhibited long regions of synteny (Figure 2b, Supplemental Figure 4). The distribution of four-fold synonymous (degenerative) third-codon transversion (4DTv) supported this lineage-specific WGD (Figure 2c). Comparison of intraspecific *dS* between homologous gene pairs in the *Diospyros* genome and interspecific *dS* between the orthologs from *Diospyros* and *Actinidia*, or from *Diospyros* and *Vitis*, indicated that the *Dd-α* event postdated the divergence of *Diospyros* and *Actinidia*, and might coincide with the divergence of the Ebenaceae family (Figure 2c-d, Supplemental Figure 4). Two other events, *Ad-α* and *Ad-β*, have been inferred by a similar analysis in the *Actinidia* genome (Huang et al., 2013) (Supplemental Figure 5) but are not detectable in the *Diospyros* genome. Thus, *Actinidia* and *Diospyros* differ by at least three lineage-specific ancestral WGD events. These occurred at a time similar to previously reported paleoduplication events in the asterids (Huang et al., 2013; Iorizzo et al., 2016; Reyes-Chin-Wo et al., 2017), as well as across the angiosperms (Vanneste et al., 2014; Van de Peer et al., 2017), concentrated around the K-Pg (Cretaceous-Paleogen) boundary (Figure 2e).

**Figure 4.**
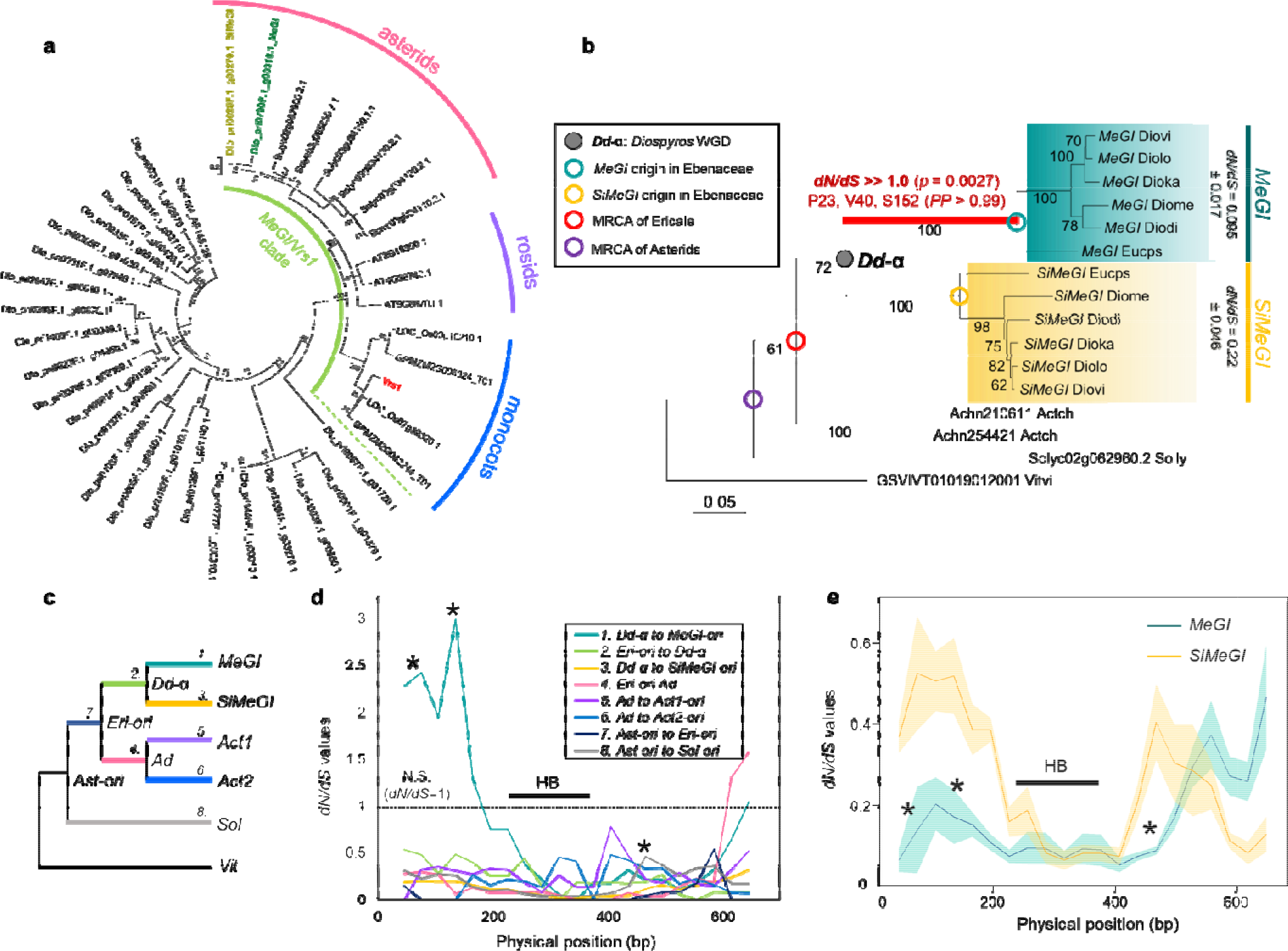
Lineage-specific adaptive evolution of *MeGI*. **a**, Phylogeny of the HD-Zip1 type homeodomain genes in the *D. lotus* genome. Only *MeGI* and *SiMeGI* were nested into the *MeGI/Vrs1*-clade with statistically significant support (100/100 and 74/100 for the divergence of *MeGI/Vrs1* clade and *MeGI/SiMeGI* subclade in *D. lotus*, respectively). **b**, Divergence of the *MeGI/SiMeGI*-like orthologs in the asterids and evidence of strong positive selection immediately after the *Dd-α* WGD event in *Diospyros* species (colored in red). No significant positive selection was detected elsewhere in this tree. Pairwise *dN/dS* values within the current *MeGI* (0.095) and *SiMeGI* (0.22) sequences suggest that both genes have been functionally fixed. **c**, Branch-specific *dN/dS* rates sliding window analysis of *MeGI/SiMeGI*-like genes from various asterid species. *MeGI* specifically exhibits positive selection in the 5’ region (~0-170bp). The three asterisks indicate the positions of the positively selected sites according to the site-branch specific detection analysis performed using PAML. The position of the homeobox domain (HB) is indicated by the thick black line. **d**, Sliding window assessment of the pairwise *dN/dS* values in the current *MeGI* and *SiMeGI* alleles. All three of the positions positively selected in *MeGI* sites after the *Dd-α* WGD event (asterisks) are under stronger purifying selection in *MeGI* than in *SiMeGI*, consistent with a situation of an adaptive evolution utilizing the mutations positively selected after WGD.

**Figure 5.**
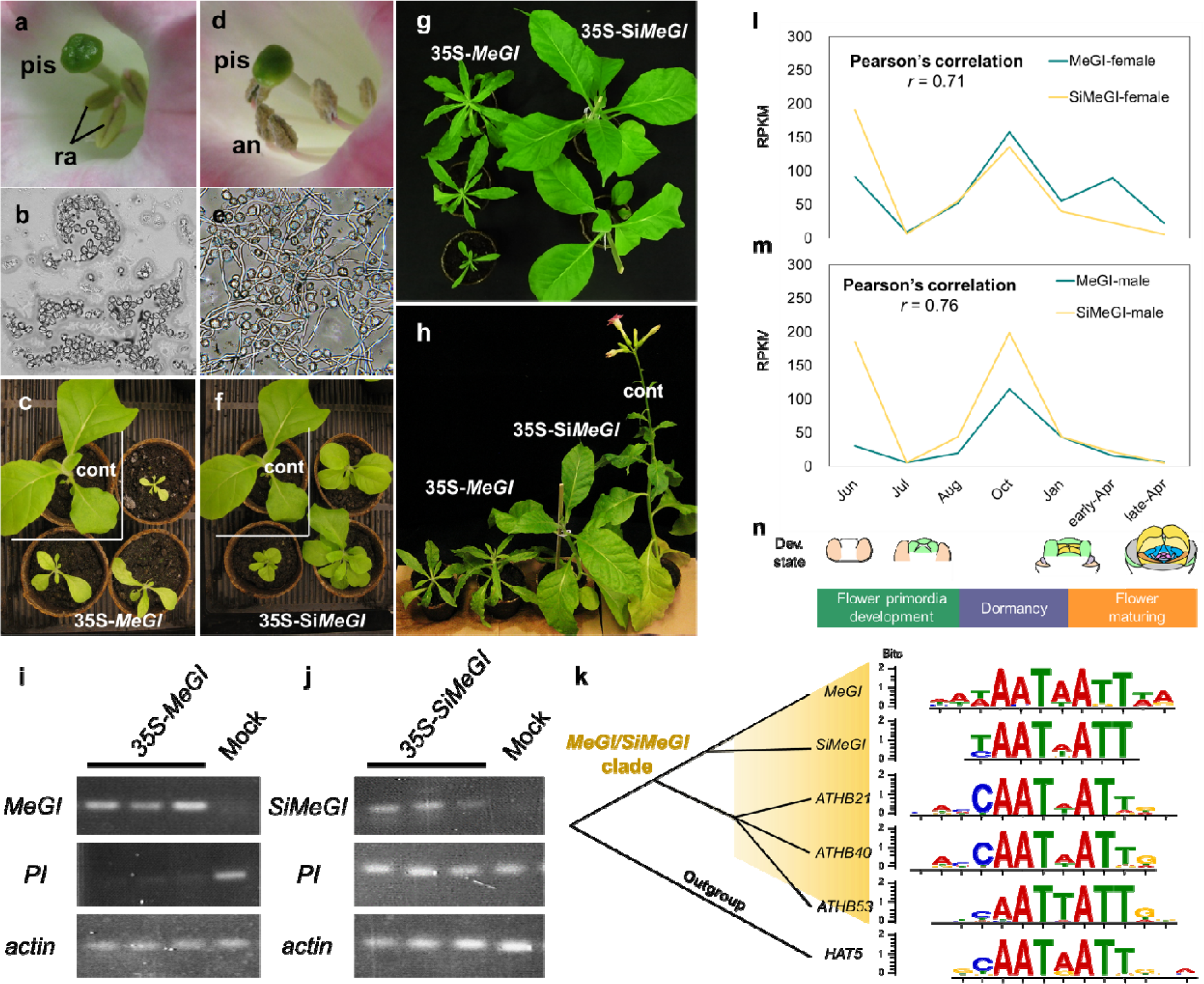
Functional differentiation between *MeGI* and *SiMeGI*. **a-h**, *N. tabacum* transgenic lines expressing either of *MeGI* or *SiMeGI* under the control of the 35S promoter. The lines expressing *MeGI* (**a-c**) showed rudimental anthers (**a**) which did not produce functional pollen grains (**b**), and severe dwarfism with chlorophyll starvation and narrow leaves (**c**, see Supplemental Figure 7 for the detail). The lines expressing *SiMeGI* (**d-f**) developed regular anthers (**d**) which produced fertile pollen (**e**), and showed moderate dwarfism (**f**). pis: pistil, ra: rudimental anthers, an: anthers. **g-h**, Both *MeGI*- and *SiMeGI*-overexpressing lines were phenotypically different from the control plants transformed with empty vectors (cont), but the *MeGI*-expressing lines exhibited more severe departure from the WT controls for specific traits, such as leaves width (see Supplemental Figure 7). Bars indicate 5mm for a and d, 50mm for c, f, g, and h. **i-j,** expression patterns of *MeGI*, *SiMeGI*, and *PI*, with *actin* as a positive control, in the transgenic lines transformed with CaMV35S-*MeGI* (i) and CaMV35S-*SiMeGI* (j). **k**, DNA motifs enriched in the DAP-Seq analyses with *MeGI* (Yang et al. 2019), *SiMeGI* (our experiments), and three Arabidopsis HD-ZIP1 genes (Khan et al. 2018) nested within the *MeGI/SiMeGI* clade. **l-n**, expression patterns of *MeGI* and *SiMeGI* in buds and flower primordia were highly correlated (Pearson’s *r* > 0.7). Expression levels in female (**l**) and male (**m**) are expressed as RPKM values. **n**, Developmental stages.

### Only a few gene, including MeGI, exhibit signs of positive selection but divergent expression patterns are common following the WGD event

To explore the evolutionary significance of lineage-specific duplications, and particularly of the *Dd-α* WGD event, *dN/dS* values between the duplicated gene pairs putatively derived from the *Dd-α* WGD events (N = 2,619) were calculated. The *dN/dS* values averaged over the coding regions indicated that most of the duplicates experienced either purifying or neutral selection (*dN/dS* ≤ 1.0, Figure 3a). In contrast, site- and evolutionary branch-specific tests for positive selection (dN/dS ≫ 1.0), using PAML, suggested that at least 9 genes experienced strong positive selection (posterior probability > 0.99 in Bayes Empirical Bayes analysis) following the *Dd-α* WGD event (Figure 3b-c). Importantly, *MeGI* and its paralog, named *Sister of MeGI* (*SiMeGI*), were one of these 9 gene pairs.

In contrast to very small number of genes exhibiting positive selection, a larger proportion of the gene pairs derived from the *Dd-α* WGD events exhibited significant differences in expression patterns. We described expression patterns in male and female buds/flowers using transcriptome data from 8 time points throughout the annual cycle (see Materials and Methods for details). Our results suggest that 45.5% of the gene pairs (597/1,311 pairs) showed significant differentiation (Pearson product-moment correlation test *r*^2^ < 0.3, Supplemental Dataset 2). To investigate differences in expression pattern between male and female flowers throughout development, we conducted 2×2 Fisher’s exact test on the *Dd-α*-derived gene pairs (see Materials and Methods) and identified 36 and 65 gene pairs (of 1,311 pairs) exhibiting significant differentiation (*p* < 0.01) in expression patterns between male and female flowers at developing and maturing stages, respectively (Figure 3d-e, Supplemental Dataset 3). These might have potentially contributed to the establishment of *Diospyros*-specific sex determining mechanisms. Such frequent variation in expression patterns is consistent with previous results in soybean (Roulin et al., 2013) and could have originated from rapid evolution in *cis*-motifs after WGD.

### Adaptive evolution of MeGI to act specifically for repression of androecium development

Genome-wide survey of the HD-Zip1 family, to which *MeGI* belongs, found 34 genes in the *D. lotus* genome. Phylogenetic analysis of *MeGI/Vrs1* orthologs from representative angiosperm species indicated that only *MeGI* and *SiMeGI* belong in the *MeGI/Vrs1* clade (bootstrap = 100/100, Figure 4a, Supplemental Figure 6). Finer evolutionary analysis on the *MeGI/SiMeGI* orthologs, to detect site-branch specific evolutionary rates using PAML, indicated that specific regions of *MeGI* experienced strong positive selection soon after the *Dd-α* event (Figure 4b, p = 0.0027 for *dN/dS* > 1.0, post. prob. > 0.99 for P23-V40-S152, and Figure 4c-d, *dN/dS* > 2.0 for the region between 45 and 165 bp in the sliding window test).

**Figure 6.**
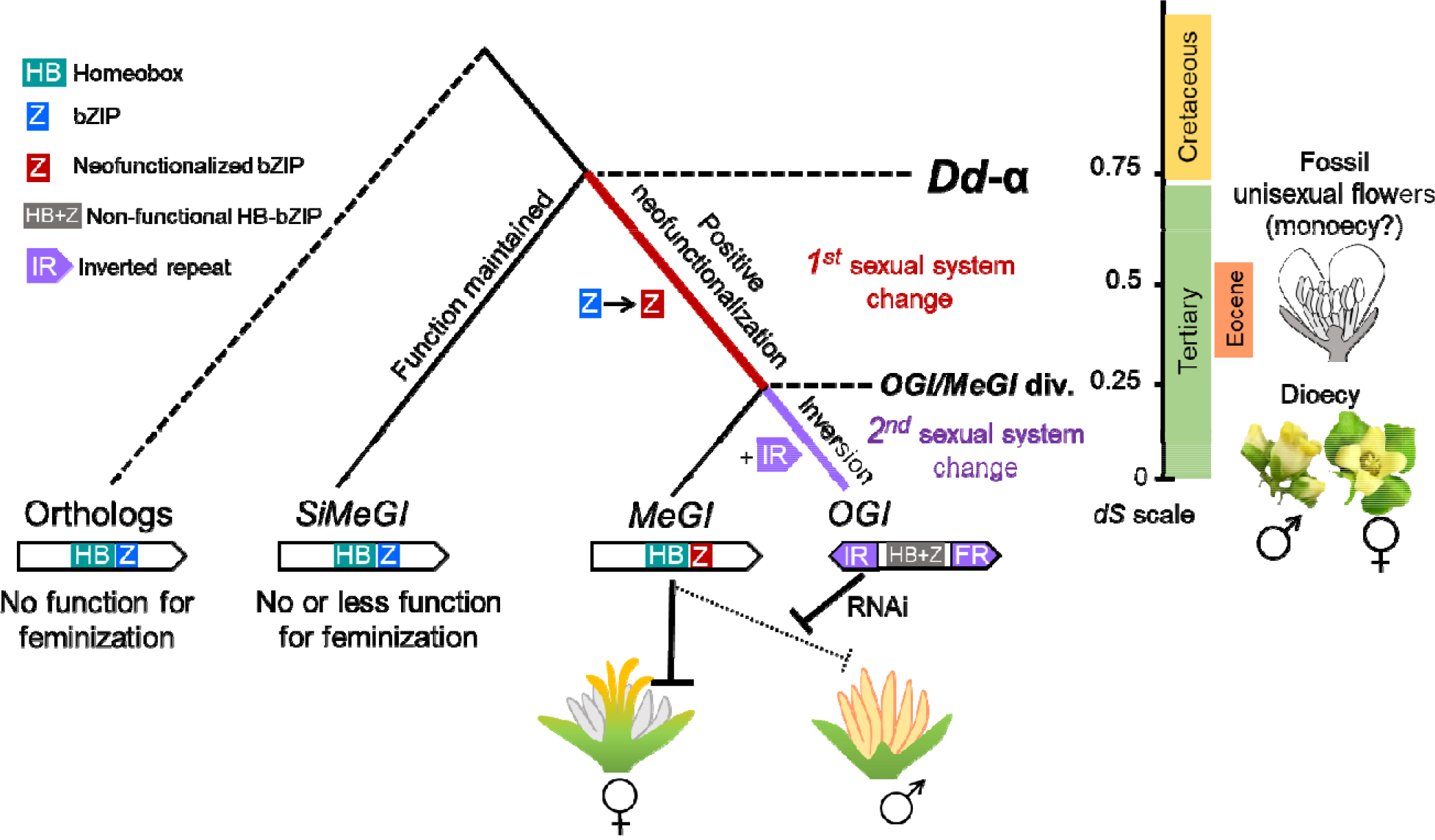
Hypothetical model for the role of duplication events in the evolution sexual systems in *Diospyros*. The *Dd-α* event triggered positive selection on the 5-end and bZIP motifs, resulting in the acquisition of a new role of *MeGI* as repressor of male organs. This was potentially associated with the first switch in sexual system, from hermaprhodistism to monoeocy. The following duplication event, a small-scale event, resulted in formation of *OGI*, containing an inverted repeat, which acquired the function of repression of *MeGI* expression via small- RNA production. This potentially triggered the establishment of the XY (heterogametic male) sexual system (Akagi et al., 2014). On the right, the *dS* scale, corresponds to the evolution of the *MeGI/SiMeGI* families and the observed sexual systems in each era. Based on the study of fossil records, unisexual (male) flowers were present during the Eocene era, which occurred significantly later than the *Dd-α* event (or the K-Pg boundary) (Christophel and Basinger, 1982). The *OGI/MeGI* divergence and the establishment of the current function of *OGI* is ancestral to diversification within *Diospyros* and consistently, the Y-encoded *OGI* regulates dioecy in the whole *Diospyros* genus (Akagi et al., 2014).

On the other hand, *MeGI* experienced strong purifying selection overall (average *dN/dS* = 0.095) since the establishment of the Ebenaceae (*Euclea* and *Diospyros*) (Figure 4b). Furthermore, the regions that experienced positive selection early are currently under stronger purifying selection in *MeGI* than in *SiMeGI* (Figure 4e). This is also consistent with the idea that *MeGI* first underwent neofunctionalization following the paleoduplication event, and that these changes were later fixed by positive selection. On the other hand, stronger purifying selection in *MeGI* than in *SiMeGI* could reflect lesser functional importance of *SiMeGI* (or possibly that it is degenerating since the duplication occurred). Alternatively, it could reflect from the need to conserve high sequence homology between *OGI* and *MeGI* in order to maintain the regulatory role of *OGI* via smRNA targeting *MeGI*.

Consistent with the evolutionary analysis presented above, ectopic expression of *MeGI* or *SiMeGI* in *Nicotiana tabacum* indicate differentiation of their protein functions. Constitutive induction of *MeGI* under the control of the CaMV35S promoter resulted in severely dwarfed plants and repressed androecium development (Figure 5a-c and g-h, Supplemental Figure 7, Supplemental Table 5), consistent with previous results using the same construct in *A. thaliana* (Akagi et al., 2014). On the other hand, constitutive induction of *SiMeGI* under the control of the same promoter resulted in plants of only slightly reduced stature and normal androecium development in *N. tabacum* (Figure 5d-h, Supplemental Figure 7, Supplemental Table 6). The function of *MeGI* as repressor of androecium in persimmon is due to the ability to regulate *PISTILLATA* (*PI*) in young developing androecium (Yang et al. 2019). The expression level of *PI* in *N. tabacum* was significantly down-regulated in the transgenic lines with *MeGI*, while the lines transformed with *SiMeGI* showed no changes in *PI* expression (Figure 5i-j). In Arabidopsis, which is a very far lineage from *Diospyros*, high expression of *SiMeGI* typically did not result in altered flower morphology although it occasionally resulted in inhibited androecium development (Supplemental Figure 8, Supplemental Tables 7 and 8).

Taken together, our results are consistent with the hypothesis that a role in androecium development is specific to *MeGI*. This is further supported by the fact that mutants of the *MeGI/SiMeGI* orthologs which are normally expressed in flower primordia in other angiosperm species, do not affect androecia development (Komatsuda et al., 2007; Whipple et al., 2011; Sakuma et al., 2013). Our evolutionary analyses revealed that the positive selection that affected *MeGI* specifically did not occur on the region binding to the target *cis*-motifs, called homeobox-domain (HB) (Figure 4d-e), but rather on the 5’ undefined region and on the leucine zipper region putatively forming heterodimers (Ariel et al., 2007; Sakuma et al., 2011). This was supported by the results of DNA affinity purification sequencing (DAP-Seq) (Bartlett et al. 2017) using *MeGI* (Yang et al. 2019) or *SiMeGI* fused to a Halo-tag, to identify which genes and/or motifs they target. The DAP-Seq reads were mapped to the *D. lotus* genome to characterize the accumulated recognition motifs (see Materials and Methods). We identified the motifs using the top 1,000 high-confidence peaks, and determined that the AATWATT sequence was enriched when using MeGI (Yang et al. 2019) and SiMeGI as the probes (Figure 5k). This motif is commonly recognized by the Arabidopsis HD-ZIP1 genes as well (Khan et al. 2018, Yang et al. 2019). Thus, it is possible that the feminization role of *MeGI* could have resulted from either increased efficiency or novel affinity to interact with other factors. Finally, the native expression patterns of *MeGI* and *SiMeGI* in persimmon are also slightly different in developing buds and flower primordia (Figure 5l-n, Supplemental Figure 9 and 10). Specifically, *MeGI* exhibits higher expression than *SiMeGI* during the flower maturing stages (Figure 5l-n). This expression differentiation might also contribute to *MeGI-*specific feminizing function.

### Transitions towards dioecy are associated with duplication events

Our results suggest the following working hypothesis for the evolutionary path into dioecy in *Diospyros*. The *Diospyros*-specific WGD event, *Dd-α* resulted in the appearance of *MeGI* and promoted the neofunctionalization of this gene into a dominant suppressor of androecium, as a feminization factor. This was followed by a second, local duplication of *MeGI* to derive a Y-encoded *OGI*, which is a dominant repressor of *MeGI* (Figure 6). Interestingly, the information available so far from other dioecious species hints at the possibility that this type of pattern may have played a role in the evolution of dioecy in other species. For example, in the establishment of dioecy in garden asparagus, the Y-encoded putative sex determinant, *SOFF*, is thought to have originated from an *Asparagus*-specific gene duplication event, which was followed by the acquisition of its function as a dominant suppressor of feminization (SuF) (Harkess et al., 2017). Furthermore, the Y-encoded putative sex determinant in kiwifruit (*Actinidia* spp.), *Shy Girl*, which acts as a dominant suppressor of feminization, also arose via an *Actinidia*-specific duplication event (Akagi et al., 2018), probably one of the *Actinidia*-specific WGD events, *Ad-α* (Huang et al., 2013). These parallel paths towards the independent evolution of all three of these sex determinants is probably not coincidental, but consistent with the theoretical framework described above. In flowering plants, transition into separated sexuality requires the appearance and selection of a gain-of-function event in order to acquire a dominant suppressor(s), such as *MeGI*. Genome-wide duplication events provide good opportunities for such a scenario. The concentration of independent paleoplodization events in the K-Pg boundary is consistent with the adaptive evolution of plants against the substantial environmental changes, including mass extinction of their pollinators that took place at the time (Wilf et al., 2006; Van de Peer et al., 2017). A selfing habit engendered by polyploidy would be advantageous, but protracted evolutionary success would be favored by an eventual return to outcrossing. The neofunctionalization of *MeGI* resulting in the acquisition of a lineage-specific new sexual system could be one of these adaptive strategies. This hypothesis is also consistent with the observed wide diversity of sex determination system within plants.

## Materials and Methods

### Initial genome sequence assembly

Dormant buds of *D. lotus* cv. Kunsenshi-male were burst in the dark for 2-weeks to harvest chlorophyll-starved young leaves. High molecular weight DNA were extracted using the Genome-tip 100/G kit (QIAGEN, Tokyo, Japan), followed by purification using phenol/chloroform extraction. Libraries were size-selected using the Blue Pippin and the following size minimums: 12 kb (14 SMRT cells), 15 kb (34 SMRT cells) and 16 kb (12 SMRT cells). A total of 60 SMRT cells and 54 Gb of PacBio raw data were obtained using the PacBio RSII. Filtered sub-reads were pooled and the longest were retained for assembly, by removing all filtered subreads shorter than 12 kb. This resulted in approximately 32x coverage of the estimated 1 Gb haploid genome size. PacBio reads were assembled using Falcon, producing 3,417 primary contigs and 6,318 alternate contigs. Next, all contigs were assessed for the presence of contaminating sequences by aligning each contig to a custom database using BLASTN+ version 2.2.31+. The custom database contained Kiwifruit psuedomolecule (ftp://bioinfo.bti.cornell.edu/pub/kiwifruit/Kiwifruit_pseudomolecule.fa.gz), the *A. thaliana* chromosomes (ftp://ftp.arabidopsis.org/home/tair/Genes/TAIR10_genome_release/TAIR10_chromosome_files/), as well as the human draft genome and representative bacterial / archaeal genome databases (pre-formatted blast+ database ftp://ftp.ncbi.nlm.nih.gov/blast/documents/blastdb.html). Hits to the two contaminant databases were identified and used to remove sequences that were largely contaminant, or to trim those with non-contaminant sequences at least 10kb long. After this step, 3,252 primary and 5,939 alternative contigs were retained. This set of contaminant-free contigs were next polished using quiver (version 2.3.0-140936) and default parameters. After this last step, 3,073 primary and 5,901 alternative contigs remained.

### Illumina library construction and sequencing

#### 1. Genomic libraries

Approximately 1.5 μg of genomic DNA was used for the construction of Illumina genomic libraries; the DNA was fragmented using NEBNext dsDNA Fragmentase (New England BioLabs; NEB) for 40–60 min at 37°C and cleaned using Agencourt AMPure XP (Beckman Coulter Genomics, Tokyo, Japan) for size selection. To select fragments ranging between 300 and 600 bp, 27 μl of AMPure was added to the 63 μl reaction. After a brief incubation at RT, 90 μl of the supernatant was transferred to a new tube and 20 μl water and 30 μl AMPure were added. After a second brief incubation at RT, the supernatant was discarded and the DNA was eluted from the beads in 20 μl of water, as recommended. Next, DNA fragments were subjected to end repair using NEB’s End Repair Module Enzyme Mix, and A-base overhangs were added with Klenow (NEB), as recommended by the manufacturer. A-base addition was followed by AMPure cleanup using 1.8:1 (v/v) AMPure reaction. Barcoded NEXTflex adaptors (Bioo Scientific, Austin, USA) were ligated at room temperature using NEB Quick Ligase (NEB) following the manufacturer’s recommendations. To remove contamination of self-ligated adapter dimers, libraries were size-selected using AMPure in 0.8:1 (v/v) AMPure:reaction volume to select for adapter-ligated DNA fragments at least 400-bp long. Half of the eluted DNA was enriched by PCR reaction using Prime STAR Max (Takara, Tokyo, Japan) at the following PCR conditions: 30 s at 98°C, 10 cycles of 10 s at 98°C, 30 s at 65°C and 30 s at 72°C and a final extension step of 5 min at 72°C. Enriched libraries were purified with AMPure (0.7:1 v/v AMPure to reaction volume), and quality and quantity were assessed using the Agilent BioAnalyzer (Agilent Technologies, Tokyo, Japan) and Qubit fluorometer (Invitrogen, Waltham, USA). Libraries were sequenced using Illumina’s HiSeq 2500 or HiSeq4000 (150-bp paired-end reads).

#### 2. GBS/ddRAD-Seq libraries

Two F1 mapping populations, derived from crosses between two *D. lotus*, Kunsenshi-male and Kunsenshi-female, and between two *D. lotus*, Kunsenshi-male and Budogaki-female, were employed for ddRAD-Seq (Peterson et al., 2012) and GBS (Elshire et al., 2011) analyses to construct genetic linkage maps. The former and latter mapping populations were named KK (n = 314) and VM (n = 119), respectively. Genomic DNA was extracted from the leaves of each line using the CTAB method. The ddRAD-Seq libraries for KK and VM were constructed using restriction enzymes *PstI* and *Msp*I (Shirasawa et al., 2016), while the GBS library for KK were prepared using with *Pst*I (Elshire et al., 2011).

#### 3. mRNA libraries

Developing buds and flowers from two *D. lotus* individuals, Kunsenshi-male and Kunsenshi-female, were harvested from June to April to cover the annual cycle of leaves/flower development. Total RNA was extracted using the Plant RNA Reagent (Invitrogen) and purified by phenol/chloroform extraction. Five micrograms of total RNA was processed in preparation for Illumina Sequencing, according to a previous report (Akagi et al., 2014). In brief, mRNA was purified using the Dynabeads mRNA purification kit (Life Technologies, Tokyo, Japan). Next, cDNA was synthesized via random priming using Superscript III (Life Technologies) followed by heat inactivation for 5 min at 65°C. Second-strand cDNA was synthesized using the second-strand buffer (200 mM Tris–HCl, pH 7.0, 22 mM MgCl_2_ and 425 mM KCl), DNA polymerase I (NEB, Ipswich, USA) and RNaseH (NEB) with incubation at 16°C for 2.5 h. Double-stranded cDNA was purified using AMPure with a 0.7:1 (v/v) AMPure to reaction volume ratio. The resulting double-stranded cDNA was subjected to fragmentation and library construction, as described above, for genomic library preparation. Ten cycles of PCR enrichment were performed using the method described above. The constructed libraries were sequenced on Illumina’s HiSeq 4000 sequencer (50-bp single-end reads).

#### 4. DAP-Seq libraries

The DAP genomic DNA libraries were prepared as previously described (O’malley et al., 2016, Bartlet et al., 2017, Yang et al. 2019). Briefly, the Covaris M220 ultrasonicator (with the manufacturer-recommended setting) was used to fragment gDNA to an average size of 200 bp. The resulting fragmented gDNA was ligated to the NEXTflex adaptors (Bioo Scientific, Austin, USA) as described, to make genomic libraries. The full-length *SiMeGI* cDNA was cloned into the pDONR221 vector (Life Technologies) and then transferred to the pIX-Halo using LR clonase II (Life Technologies) to generate pIX-Halo-SiMeGI. pIX-Halo-MeGI has been constructed previously (Yang et al. 2019). The N-terminally Halo-tagged *MeGI* and *SiMeGI* were produced using the TNT SP6 Coupled Wheat Germ Extract System (Promega, Fitchburg, WI, USA) and purified with Magne HaloTag beads (Promega). A total of 50 ng DAP gDNA library was incubated with Halo-tagged *MeGI* and *SiMeGI* at room temperature for 1 h.

#### 5. Sequencing

The ddRAD Seq sequences were obtained at the Kazusa DNA Research Institute. The GBS sequences were obtained from the Genomic Diversity Facility (Cornell University). All other Illumina sequencing were conducted at the Vincent J. Coates Genomics Sequencing Laboratory at UC Berkeley, and the raw sequencing reads were processed using custom Python scripts developed in the Comai laboratory and available online (http://comailab.genomecenter.ucdavis.edu/index.php/Barcoded_data_preparation_tools), as previously described. In brief, reads were split based on index information and trimmed for quality (average Phred sequence quality > 20 over a 5 bp sliding window) and adaptor sequence contamination. A read length cut-off of 35 bps was applied to mRNA reads. Sequencing analysis of ddRAD-Seq libraries was performed at the Kazusa DNA Research Institute, and data processing was conducted as described in Shirasawa et al., (2016). All samples used to generate Illumina sequences are listed in Supplemental Table 10.

### Gene prediction and genome/genes annotation

The RNA-Seq data for gene prediction was obtained from developing buds and flowers from *D. lotus* Kunsenshi-male at the following eight time points in 2013 to 2015 (June, July, August, October, January, March, early April, and late April) to cover the annual cycle of leaves/flower development. The RNA-Seq reads were trimmed according to previous reports (Akagi et al., 2014). The cleaned reads were mapped onto the scaffolds of DLO_r1.1 using TopHat 2.0.14 (Trapnell et al., 2009), and the BAM files obtained were used for BRAKER1 1.9 pipeline (Hoff et al., 2016). In the pipeline, GeneMark-ET 4.32 (Lomsadze et al., 2018) and Augustus 3.1 (Stanke and Waack, 2003) were used to construct the training set, and Augustus 3.1 was used for the gene prediction, using the training set. Genes were compared to the UniProtKB (http://www.uniprot.org/uniprot/) and of Araport11 (Krishnakumar et al., 2015) peptide sequences using BLASTP with E-value cutoff of 1E-10. Genes that were similar to those in the databases were categorized as “highly confident” (HC). Analysis of the conservation of the single-copy genes was conducted using BUSCO v1 (Simão et al., 2015). Repeat sequences were detected using RepeatScout 1.0.5 (Price et al., 2005) and RepeatMasker 4.0.6 (http://www.repeatmasker.org) against the Repbase database (Bao et al., 2015), according to the method used previously (Hirakawa et al., 2014). The HC genes on the primary scaffolds (DLO_r1.1 primary) were compared to the genes of *Actinidia chinensis* (kiwifruit; 39,040 genes (Huang et al., 2013)), *Vitis vinifera* (grape; 29,927 genes (IGGP 12x.31) (Jaillon et al., 2009)), *Solanum lycopersicum* (tomato; 34,789 genes (ITAG 3.10) (The Tomato Genome Consortium, 2012)) and *Arabidopsis thaliana* (27,655 genes (Araport11)) using OrthoMCL 2.0.9. To estimate the divergence time between *D. lotus*, *A. chinensis*, *V. vinifera*, and *A. thaliana*, the single copy genes conserved amongst all four species were aligned by MUSCLE 3.8.31 (Edgar, 2004). InDels in the alignment were eliminated using Gblocks 0.91b (Castresana, 2000), and the sequences were concatenated by species and used to construct the phylogenetic tree using the Maximum Likelihood method using MEGA 7.0.26 (Tamura et al., 2013) with the Jones-Taylor-Thornton (JTT) model as the substitution model. The divergence time was estimated based on that between *A. chinensis* and *V. vinifera* (117 MYA) published in TIMETREE (http://www.timetree.org).

### Construction of the persimmon database

The sequence data obtained was released in the form of the PersimmonDB (http://persimmon.kazusa.or.jp). In the database, BLAST searches can be conducted against the scaffolds (DLO_r1.0) and pseudomolecules (DLO_r1.0_pseudomolecules), cds (DLO_r1.1_cds), and pep (DLO_r1.1_pep). Keyword searches are available against the results of the similarity searches against TrEMBL and peptide sequences in Araport11. The genomic and genic sequences, GFF files of the scaffolds and pseudomolecules, and BED files can be downloaded from the database. The scaffolds are also available under accession numbers BEWH01000001-BEWH01008974 (8,974 entries) in DDBJ. The raw sequence data is also available from under accession numbers DRA006168 (Illumina WGS for *D. lotus* Kunsenshi-male and female), DRA006169-DRA006176 (ddRAD-Seq/GBS for KK and VM populations), DRA006177 (RNA-Seq for *D. lotus* Kunsenshi-male), and DRA006182-DRA006184 (PacBio WGS for *D. lotus* Kunsenshi-male) in DDBJ.

### Genetic anchoring of the scaffold using two mapping populations

The sequence reads from the ddRAD-Seq and GBS libraries were mapped onto the primary contigs of the DLO_r1.0 reference sequence using Bowtie 2 (version 2.2.3) (Langmead and Salzberg, 2012). SNP calling was performed using the mpileup command of SAMtools (version 0.1.19) (Li et al., 2009) and the view command of BCFtools (Li et al., 2009). High-confidence SNPs were selected using VCFtools (version 0.1.12b) (Danecek et al., 2011) using the following parameters: ≥10×◻coverage of each sample (--minDP 10);◻>999 SNP quality value (--minQ 999);◻≥0.2 minor allele frequency (--maf 0.2), and <0.5 missing data rate (--max-missing 0.5). Totals of 3,535 and 4,027 high-confident SNPs were obtained in the KK and VM populations, respectively. Genotype information for all lines were prepared for the CP mode of JoinMap (version 4) and classified into groups using the Grouping Module of JoinMap with LOD scores of 4 to 7. Marker order and relative map distances were calculated using its regression-mapping algorithm with the following parameters: Haldane’s mapping function ≤0.35 recombination frequency, and ◻≥2.0 LOD score. LPmerge (version 1.5) (Endelman et al., 2014) was used to integrate the linkage maps into a single consensus map. To construct pseudomolecule sequences, scaffolds assigned to the genetic map for the Kunsenshi-male, the cultivar used for the genome sequencing analysis, were ordered and oriented in accordance with marker order if at least two marker loci were mapped on a single scaffold. Otherwise, in the cases of a single marker on a scaffold, the orientation of the sequence was determined as “unknown”.

### Comparative genomics

Whole genome-resequencing analysis on the Kunsenshi-male and female individuals were performed as described in Shirasawa et al. (2017). Paired-end sequences reads were obtained from the male and female lines with Illumina NextSeq, and trimmed and filtered based on quality score using Prinseq (Schmieder and Edwards, 2011) and base similarity to adapter sequences, AGATCGGAAGAGC, using fastx_clipper in the FASTX-Toolkit (http://hannonlab.cshl.edu/fastx_toolkit). The resulting reads were mapped on the primary contigs of DLO_r1.0 reference sequence with Bowtie2, and single nucleotide polymorphisms were detected with SAMtools mpileup (Li et al., 2009) and filtered with the conditions of sequence depth of ≥10 in each line (--minDP 10) and mapping quality of >200 in each SNP locus (--minQ 200) using VCFtools (Danecek 2011). The effect of SNPs on gene function were predicted with SnpEff (Cingolani et al., 2012) to assign the SNPs to four impact categories, high, moderate, modifier, and low, predifined by SnpEff. Synteny relationship of the genome structures were predicted with PROmer program of Mummer package (Delcher et al., 2002) between *Diospyros* (this study) and *Actinidia* (Huang et al., 2013), as well as within the *Diospyros* genus. The results were filtered with delta-filter (parameters of AAA and BBB), and visualized using Circos (Krzywinski et al., 2009).

### Detection of genetic diversity within paralogs

Genes annotated as potential transposable elements by blastn/blastp using the TAIR/nr databases, and potentially repetitive genes which produced >5 homologous genes in the *D. lotus* genome (<e^−20^ in blastp), were discarded. Gene pairs showing significant sequence similarity (<e^−20^ in blastp) and their orthologs from three species, *Actinidia*, *Solanum* and *Vitis* were subjected to in-codon frame alignment using their protein and nucleotide sequences with Pal2Nal and MAFFT ver. 7 under the L-INS-i model. The resulting alignments were subjected to Mega v.6 to estimate the Jukes and Cantor corrected values of synonymous (*dS*) and non-synonymous (*dN*) substitutions and the index of evolutionary rate (*dN/dS*). The four-fold degenerative sites were extracted from the alignments with PAML (icode=11), and their pairwise transversion rates (4DTv) were calculated according to previous reports (Tuskan et al., 2006). To estimate the divergence time between the gene pairs, we adopted an estimated rate of 2.81 × 10^−9^ substitutions per synonymous site per year, according to the report in *Actinidia* (Shi et al., 2010).

### Evolutionary analysis on the paralogs derived from *Dd-α* WGD event

To search for signs of positive selection, aligned nucleotide sequences of each gene pair and an outgroup ortholog, from either the *Actinidia*, *Solunum* or *Vitis* genomes, were subjected to codon-based detection of positive selection test using PAML (Yang, 1997). The statistical significance of positive selection on branches was evaluated using the likelihood ratio test of the null hypothesis that *dN/dS* = 1. Site-specific positive selection was assessed by Bayes Empirical Bayes analysis. To examine the positively selected sites common across the all three outgroups, in-frame alignments of the *D. lotus* gene pairs with the orthologs from all of the *Actinidia*, *Solanum* or *Vitis* genomes were used for the construction of evolutionary topologies using ML method by Mega v. 6, using the general time reversible (+I+G) model. Based on these alignments and topology, the branch- and site-specific positive selection test was performed using PAML, as well.

To define the phylogenetic relationship between the *MeGI*/*SiMeGI*-like orthologs/paralogs in angiosperms, genes showing significant homology (<1e^−10^ in blastp analysis) to a HD-ZIP1 OsHOX4 from *Oryza sativa*, which was previously used as the outgroup gene for the *MeGI* clade (Akagi et al. 2014), were collected from the *Diospyros lotus*, *Solanum lycopersicum, Arabidopsis thaliana, Oryza sativa*, and *Zea mays* genomes. A total of 174 protein sequences from these genomes, and that of Vrs1 from barley (Komatsuda et al. 2007) were aligned using MAFFT ver. 7, followed by manual pruning with SeaView. The pruned alignment was subjected to the NJ approach using Mega v. 6, with the JTT model, to construct phylogenetic tree (Supplemental Figure 6).

To assess selective pressure on *MeGI* and *SiMeGI*, their alleles from other members of the Ebenaceae family (*Diospyros* and *Euclea* genera), and their orthologs in the *Actinidia*, *Solunum* or *Vitis* genomes were subjected to in-codon frame alignment by MAFFT ver. 7, followed by a ML approach using Mega v. 6, with HKY+G model, to construct an evolutionary topology. The putative ancestral sequences of the *MeGI* and *SiMeGI* origins in the Ebenaceae family, and the sequences in the most recent common ancestor (MRCA) of the order Ericale and of the Asterids, were estimated using Mega. Informative SNPs in the aligned sequences were analyzed by DnaSP 5.1 (Librado and Rozas, 2009) and used to calculate a series of window-average *dN/dS* values, from the start codon (ATG) in a 150-bp window with a 30-bp step size, until the walking window reached the stop codon. To assess differentiation of expression patterns between the *Dd-α*-derived paralog pairs, we conducted Pearson’s product moment correlation analysis and Fisher’s exact test. Differentiation between the developmental stages of the buds/flowers throughout the annual cycle was examined by the “cor.test” function in R (with “pearson” method), using mRNA-Seq transcriptome data from datapoints (Supplemental Dataset 3). Differentiation of expression pattern between male and female flowers was examined for each paralog pair using a 2×2 Fisher’s exact test (“fisher.test” function in R), and using mRNA-Seq transcriptome data from early developing stage and maturing stage, respectively (Supplemental Dataset 3).

### Transformation of *MeGI* and *SiMeGI*

Full length sequences of the *MeGI* and *SiMeGI* transcripts were amplified by PCR using PrimeSTAR Max (TaKaRa) from cDNA synthesized from RNA, itself derived from developing flower buds of *D. lotus* cv. Kunsenshi-male (Supplemental Tables 10 and 11). The amplicons were cloned into the pGWB2 vector to place the genes under the control of CaMV35S promoter. We constructed pGWB2-*MeGI* and pGWB2-*SiMeGI* using the Gateway system (Invitrogen) and the pENTR/D-TOPO cloning kit and LR clonase. Tobacco plants (*N. tabacum*) cv. Petit Havana SR1 were grown *in vitro* under white light with 16-h-light and 8-h-dark cycles at 22°C until transformation. The binary construct was introduced into the *A. tumefaciens* strain EHA101. Young petioles and leaves of tobacco plants were transformed by the leaf disk method as previously described (Akagi et al., 2014). Transgenic plants were selected on Murashige and Skoog medium supplemented with 100 μg/mL kanamycin. Pollen tube germination was assessed 6 h after placing the pollen grains on 15% sucrose/0.005% boric acid/1.0% agarose media at 25°C. The pollen germination ratio was counted as average percentages in batches of 200 pollen grains from the first three flowers.

### RNA *in situ* hybridization

RNA *in situ* hybridization was performed as previously described (Esumi et al., 2007), but with minor modifications. Briefly, bud samples were fixed in FAA (1.8% formaldehyde, 5% acetic acid, 50% ethanol), dehydrated using an ethanol: t-butanol series, and then embedded in paraffin. The embedded tissues were sliced into ca 10-μm sections, and the sections were mounted on FRONTIER coated glass slide (Matsunami Glass Ind., Japan). Paraffin was removed with xylene, and the tissue sections were rehydrated in an ethanol series. The tissue sections were then incubated in a Proteinase K solution (700U/mL Proteinase K, 50mM EDTA, 0.1M Tris-HCl pH 7.5) for 30 min at 37°C, followed by acetylation with acetic anhydride (0.25% acetic anhydride in 0.1 M triethanolamine solution) for 10 min. Full length *MeGI* and *SiMeGI* cDNA sequences were cloned into the pGEM-T Easy vector (Promega, WI, USA) to synthesize the DIG-labelled probes, respectively. Antisense RNA probes were synthesized using the DIG-labeling RNA synthesis kit (Roche, Switzerland), according to the manufacturer’s instruction. The probe solution including RNaseOUT (Thermo Fisher Scientific, Waltham, USA) was applied to the slides and covered with parafilm. Hybridization was performed at 48°C for >16 h. For detection, 0.1% Anti-Digoxigenin-AP Fab fragments (Sigma-Aldrich, St. Louis, USA) was used as the secondary antibody to stain with NBT/BCIP solutions.

## Author contributions

T.A., I.M.H. and L.C. conceived the study, T.A. and I.M.H., preponderantly designed the experiments. T.A. and K.S. performed the experiments. T.A., K.S., H.N., H.H. and I.M.H. analyzed the data. T.A. and R.T. initiated and maintained the plant materials. T.A., I.M.H. and L.C. drafted the manuscript. All authors participated in data interpretation, edited the manuscript and approved the final manuscript.

## Acknowledgements

We thank Dr Deborah Charlesworth for the extensive discussions on the interpretation of our results and the many thoughtful pieces of advises provided through this work. We thank Meric Lieberman (UC Davis Genome Center) for bioinformatics support, Ayaka Sugimoto and Yang Ho-Wen (Graduate School of Agriculture, Kyoto University) for experimental support. Some of this work was performed at the Vincent J. Coates Genomics Sequencing Laboratory at UC Berkeley, supported by NIH S10 OD018174 Instrumentation Grant. This work was supported by PRESTO, Japan Science and Technology Agency (to TA), and Grant-in-Aid for Young Scientists (A) (no. 26712005 to TA), for Challenging Exploratory Research (no. 15K14654 to TA), Grant-in-Aid for Scientific Research on Innovative Areas No. J16H06471 to TA from JSPS, and by the National Science Foundation (NSF) IOS award under Grant No. 1457230 (to IMH and LC).

## Supplementary Information

**Supplemental Figure 1: kmer distribution to estimate genome size and the degree of heterozygosity.**

The distribution of distinct *k*-mers (k◻=◻17) from the Illumina short reads showed two peaks at multiplicities of 32 and 54. The low and high peaks represent heterozygous and homozygous sequences, respectively. We estimated the genome size to be 877.7◻Mb from the higher peak. This estimation almost agreed with the value measured by flow cytometry, 907 Mb, which was calculated from the nuclear DNA content in *D. lotus* of 1.85◻pg/2◻C (Tamura et al., 1998) and an assumption that 1◻pg of DNA is equivalent to 980◻Mb (Bennett et al., 2000).

**Supplemental Figure 2: Genetic anchoring data (Dlo01-15).**

Genetic linkage map (left bars) and physical map of Dlo_r1.0 pseudomolecule sequences (right bars). Colors in the genetic map represent density of SNPs per 5 cM, while black, white and gray bars in the physical map indicate the positions of the forward, invert, and unknown directional scaffold sequences integrated in the pseudomolecule, respectively.

**Supplemental Figure 3: Conservation of gene and repetitive sequences across representative plant species**

**a**, Amino acid sequences were compared among genes from *D. lotus* (40,532 genes; DLO_r1.1 primary), *A. chinensis* (39,040 genes (Huang et al., 2013)), *V. vinifera* (29,927 genes (IGGP 12x.31) (Jaillon et al., 2009)), *S. lycopersicum* (34,789 genes (ITAG 3.10) (The Tomato Genome Consortium, 2012)), and *A. thaliana* (27,655 genes (Araport11) (Cheng et al., 2017)) using OrthoMCL v2.0.9 (Li et al., 2003) with default parameters. The numbers of clusters were shown in the intersections of the Venn diagram. **b,** Repetitive sequences were identified by RepeatMasker v4.0.6 (http://www.repeatmasker.org) using Repbase v406 (http://www.girinst.org/repbase/) and RepeatScout v1.0.5 for the genome sequences of *D. lotus* (8,974 sequences; DLO_r1.0), *A. chinensis* (30 pseudomolecules (Huang et al., 2013)), *S. lycopersicum* (13 pseudomolecules (SL3.0) (The Tomato Genome Consortium, 2012)), *Lactuca sativa* (lettuce; 9 pseudomolecules (V8) (Reyes-Chin-Wo et al., 2017)), *V. vinifera* (33 chromosomes (IGGP 12x.31) (Jaillon et al., 2009)), *Prunus persica* (peach; 8 pseudomolecules (v2.0.a1) (Verde et al., 2013), *Carica papaya* (papaya; 5,901 scaffold sequences (ASGPBv0.4) (Ming et al., 2008)), and *A. thaliana* (5 chromosomes, chloroplast and mitochondria genomes (TAIR10)). The percentage of repetitive sequences against the total length of the genome sequence were calculated for each of the result of RepeatMasker and RepeatScout and compared among the plant species.

**Supplemental Figure 4: Gene duplication patterns following the *Dd-α* event.**

**a**, Heat map for the numbers of genes derived from *Dd-α*, shared between two chromosomes. For instance, Dlo01 shared many paralogs with Dlo02 and Dlo12, while Dlo02 and Dlo12 shared few paralogs with each other. The patterns of such affinities between the chromosomes suggests a paleotetraploidization event. **b,** Syntenic relationship betweem the putatively *Dd-α*-derived paralogous genes within the *Diospyros* genome. **c**, Synteny dot plot in a putative paleoduplicated region, on chromosomes 1 and 2. The red dotted boxes indicate long segmental syntenic blocks (>3Mb).

**Supplemental Figure 5: Genome-wide synteny between *Diospyros* and *Actinidia***

Dot plots of the syntenic genomic regions between *Diospyros* and *Actinidia*. As represented in the orange box, a single genomic segment from *Actinidia* corresponds to two syntenic *Diospyros* genome regions which are derived from the *Dd-α*. In the orange box, the middle regions of Dlo01 and Dlo02, and Dlo03 and Dlo06 are duplicated regions via *Dd-α* (see Figure 1 and Supplemental Figure 4). On the other hand, as represented in the green box, a genomic segment from *Diospyros* corresponds to at maximum four syntenic *Actinidia* genome regions which are derived from the double *Actinidia*-specific paleoduplication events (Ad-α and Ad-β) (Huang et al., 2013). These results indicate that, in the evolution of the order Ericales, *Dd-α* and Ad-α/β occurred independently in the *Diospyros* and *Actinidia* ancestral genomes, respectively.

**Supplemental Figure 6: Phylogenetic tree of the angiosperm HD-ZIP1 family.**

Phylogeny of the HD-Zip1 type homeodomain genes in representative angiosperm genomes (*Solanum lycopersicum, Oryza sativa, Zea mays*, and *Arabidopsis thaliana*) and the *D. lotus* genome. The 175 HD-ZIP1 genes clustered into 8 major clades (Clade I-VIII), which each included at least one homologs from all 5 species used in this study (see Materials and Methods), although the root branches of clades VI and VII were not statistically significant (32/100 and 28/100, respectively). *MeGI*, *SiMeGI* and Vrs1 were nested within clade IV (colored in light green).

**Supplemental Figure 7: Overexpression of *MeGI* and *SiMeGI* under the control of CaMV35S promoter in *N. tabacum***

**a-c**, 1-week old transgenic lines. The *MeGI*-induced lines (**a**) frequently showed clear irregularities in development, in comparison to the *SiMeGI* lines (**b**) or empty cassette-induced lines (**c**). **d**, Comparison of 4-weeks old transgenic plants. The *MeGI*-induced lines (center) uniformly showed more severe growth inhibition, than the *SiMeGI*-induced lines (left). **e**, close-up picture of the *MeGI*-induced line corresponding to the individual marked with an asterisk in the panel (**d**). The leaves showed irregular shapes with significantly less veins. **f**, comparison of the appearance of 15-weeks old transgenic lines. The *MeGI*- induced line (left) exhibited dwarfism, but the total number of leaves were comparable to the control plants (right), while the internode lengths were shorter than the control, as shown in the panel “**g**”. **h**, Differentiation of the leave shapes and structures in the control (left) and the *MeGI*-induced line (right). The *MeGI*-induced lines produced narrow and serrated leaves. Bars indicate 10mm for a-c, and e; 50mm for d, f, g, and h.

**Supplemental Figure 8: Overexpression of *MeGI* and *SiMeGI* under the control of CaMV35S promoter in *A. thaliana.***

**a**, Dissection of the control Arabidopsis plant transformed with an empty cassette. an: anther, pe: petal, sg: stigma. **b-e**, p35S-*MeGI* transgenic lines. Dissected flowers show rudimental anthers (ra) (**b-c**). Approximately half of the transgenic plants are semi-dwarf (semi-dwf) (**d**) or complete dwarf (**e**). They also frequently showed leaf serration, which is consistent with our previous analysis of the p35S-*MeGI* induced Arabidopsis plants (Akagi et al., 2014). **f-i**, p35S-*SiMeGI* transgenic lines. The transgenic plants occasionally showed rudimental anthers similar to the *MeGI*-induced lines (**f-g**). A part of the *SiMeGI*-induced lines showed semi-dwarfism (**h**), but full dwarfism was never observed in the 63 transgenic lines. Over 95% of the *SiMeGI*-induced lines were hermaphroditic, where the numbers of stamens are properly maintained (**i**), in contrast to the *MeGI*-induced lines (Akagi et al., 2014). Bars indicate 1mm for a, b, f, and i; 0.1mm for c and g; 10mm for d-e and h. j, Distribution of the number of female (fe) and hermaphrodite (herm) individuals in the p35- *MeGI* (green), p35S-*SiMeGI* (yellow), and p35S-empty (cont; gray) transgenic lines. k, Distribution of the number of complete dwarf (dwf), semi-dwarf (semi-dwf) and normal individuals in the p35-*MeGI*, p35S-*SiMeGI*, and p35S-empty (cont) transgenic lines.

**Supplemental Figure 9: *in situ* RNA hybridization**

RNA *in situ* hybridization in developing buds and flower primordia, using *MeGI* (**a-c**) and *SiMeGI* (**d-f**) sequences as probes. In the cross section of the developing buds (**a** and **d**), the *MeGI* signal is strong in flower buds only (fb) (**a**), while *SiMeGI* showed significant signal in the pith (Pi) and young leaves (ly), as well as in flower buds (**d**). This is consistent with our expression analyses using laser capture micro-dissected (LCM) samples (Supplemental Figure 10). In the longitudinal sections of the developing buds (**b** and **e**), both *MeGI* and *SiMeGI* signals are confined to the meristematic region, especially in the shoot apical meristems (sam). At a later developing stage (**c** and **f**), flower primordia (fp) and bract (br) showed substantial signals of both *MeGI* (**c**) and *SiMeGI* (**f**). Bars indicate 50μm.

**Supplemental Figure 10: Expression analysis of *MeGI/SiMeGI* in laser capture microdissection (LCM)**

Longitudinal (**a**) and cross (**b**) sections of buds from *D. lotus*, Kunsenshi-male, and the target of the LCM. We targeted flower buds (red), young leaf or leaf buds (blue), and pith or cambium (green). **c**, the section after laser captions. **d**, qRT-PCR analysis to detect relative expression of the *MeGI* and *SiMeGI* among the organs, at early developmental stages (Jun-Jul) when the flower primordia form. Consistent with the results of the in situ hybridization (Supplemental Figure 9), *MeGI* expression was much stronger in flower buds than in pith or young leaves, while the difference in expression levels between the three organs was less drastic for *SiMeGI*. For both graphs, the expression level in flower buds was defined as “1”. **e**, comparison of the expression level of *MeGI* and *SiMeGI* in the developing flower buds. Illumina mRNA-Seq analysis was conducted on the LCM samples to detect RPKM values of *MeGI* and *SiMeGI*. *SiMeGI* was expressed higher than or comparative to the *MeGI*, in the developing flower buds. Notwithstanding, the reduction in *MeGI* expression in this stage affect the flower sexuality and the inflorescent structure (Akagi et al., 2014). **f**, relative expression of the *MeGI* and *SiMeGI* in different organs, during dormancy stage (Dec) when the development of flower primordia halt. Flower buds showed no significant expression of either *MeGI* or *SiMeGI*.

**Supplemental Table 1: Summary statistics for the initial genome assembly of *D. lotus* cv. Kunsenshi-male**

**Supplemental Table 2: Number of SNPs and length of genetic linkage maps in *D. lotus***

Genetic maps for the four parental lines of the two mapping populations (KK and VM), were built using the pseudo-test cross method. All linkage groups were anchored to the 15 chromosomes of the *D. lotus* draft genome assembly. The sex-determinant locus was mapped to linkage group 15, suggesting the Dlo15 is the sex chromosome. More detailed information about the maps and SNPs are available from the Persimmon Genome Database (http://persimmon.kazusa.or.jp)

**Supplemental Table 3: Number of annotated SNPs and indels between the female and male lines of *D. lotus* ‘Kunsenshi’**

SNPs and indels were identified from whole-genome resequencing analysis of female and male lines of *D. lotus*, and functionally annotated and classified into four categories predefined by SnpEff (Cingolani et al., 2012): high (e.g. nonsense mutations and frameshift mutations)-, moderate (e.g. missense mutations)-, modifier (e.g. intron and intergenic mutations)- and low-impact (e.g. synonymous mutations) mutations (see http://snpeff.sourceforge.net for details). Further details about these SNPs and Indels are available from the Persimmon Genome Database (http://persimmon.kazusa.or.jp).

**Supplemental Table 4: Comparison of the repeat sequences in representative eudicot genomes**

Repetitive sequences amounted for 630.2 Mb (66.6%) of the total length of the final genome assembly. Unique repeats were abundant in the *D. lotus* genomes, constituting 49.8% of all repeats. Of the known types of repeats, Class I LTR elements were observed most frequently (11.2%).

**Supplemental Table 5: Phenotypic characterization of the p35S-*MeGI N. tabacum* transformed lines.**

**Supplemental Table 6: Phenotypic characterization of the p35S-*SiMeGI N. tabacum* transformed lines.**

**Supplemental Table 7: Phenotypic characterization of the p35S-*MeGI A. thaliana* transformed lines.**

**Supplemental Table 8: Phenotypic characterization of the p35S-*SiMeGI A. thaliana* transformed lines.**

**Supplemental Table 9: Plant materials**

**Supplemental Dataset 1: Table of predicted gene locations based on BLAST results**

**Supplemental Dataset 2: Result of the Pearson correlation test for correlation between the expression patterns of paralog pairs.** For each paralog pair, the r and t-test p-values are indicated, in addition to the RPKM values at each of the 16 expression time points selected.

**Supplemental Dataset 3: Results of the Fisher Exact test** of the relationship between expression of the two paralogs in each paralog pair in male and female developing flowers. For each paralog pair, the ratio of male to female expression is indicated as well as the result of the Fisher 2 × 2 Exact test p-value.

## References

Akagi T, Henry IM, Tao R, Comai L (2014) A Y-chromosome-encoded small RNA acts as a sex determinant in persimmons. Science 346: 646–650.

Akagi T, Henry IM, Ohtani H, Morimoto T, Beppu K, Kataoka I, Tao R (2018) A Y-encoded suppressor of feminization arose via lineage-specific duplication of a cytokinin response regulator in kiwifruit. Plant Cell 30: 780–795.

Allen AM, Hiscock SJ (2008) Evolution and phylogeny of self-incompatibility systems in Angiosperms. p. 73–102. In: V. E. Franklin-Tong (ed.). Self-incompatibility in flowering plants. Springer-Verlag, Berlin.

Ariel FD, Manavella PA, Dezar CA, Chan RL (2007) The true story of the HD-Zip family. Trends Plant Sci 12: 419–426.

Bao W et al (2015) Repbase Update, a database of repetitive elements in eukaryotic genomes. Mobile DNA 6: 11.

Barrett BC (2002) The evolution of plant sexual diversity. Nat Rev Genet 3: 274–284.

Bartlett, A., O’Malley, R.C., Huang, S.C., Galli, M., Nery, J.R., Gallavotti, A. and Ecker, J.R. (2017) Mapping genome-wide transcription-factor binding sites using DAP-seq. Nat. Protoc. 1659–1672.

Charlesworth B, Charlesworth D (1978) A model for the evolution of dioecy and gynodioecy. Amer Nat 112: 975–997.

Charlesworth D, Charlesworth B (1978) Population genetics of partial male-sterility and the evolution of monoecy and dioecy. Heredity 41: 137–153.

Castresana J (2000) Selection of conserved blocks from multiple alignments for their use in phylogenetic analysis. Mol Biol Evol 17: 540–552.

Christophel DC, Basinger JF (1982) Earliest floral evidence for the Ebenaceae in Australia. Nature 296: 439–441.

Cingolani P et al (2012) A program for annotating and predicting the effects of single nucleotide polymorphisms, SnpEff: SNPs in the genome of *Drosophila melanogaster* strain *w*^1118^; *iso*-2; *iso*-3. Fly 6: 80–92.

Danecek P et al (2011) The variant call format and VCFtools, Bioinformatics 27: 2156–2158.

Delcher AL et al (2002) Fast algorithms for large-scale genome alignment and comparison. Nucl Acids Res 30: 2478–83.

Edgar RC (2004) MUSCLE: a multiple sequence alignment method with reduced time and space complexity. BMC Bioinfo 5: 113.

Elshire RJ et al (2011) A robust, simple genotyping-by-sequencing (GBS) approach for high diversity species. PLoS ONE 6: e19379.

Endelman JB, Plomion C (2014) LPmerge: an R package for merging genetic maps by linear programming. Bioinformatics 30: 1623–1624.

Esumi T et al (2007) Relationship between floral development and transcription levels of *LEAFY* and *TERMINAL FLOWER 1* homologs in Japanese pear (*Pyrus pyrifolia* Nakai) and quince (*Cydonia oblonga* Mill.). J Jpn Soc Hortic Sci 76: 294–304.

Flagel LE, Wendel JF (2009) Gene duplication and evolutionary novelty in plants. New Phytol. 183: 557–564.

Fraser LG, Tsang GK, Datson PM, De Silva HN, Harvey CF, Gill GP, Crowhurst RN, McNeilage MA (2009) A gene-rich linkage map in the dioecious species *Actinidia chinensis* (kiwifruit) reveals putative X/Y sex-determining chromosomes. BMC Genom 10: 102.

Harkess A et al (2017) The asparagus genome sheds light on the origin and evolution of a young Y chromosome. Nat Comm 8: 1279.

Hirakawa H et al (2014) Draft genome sequence of eggplant (*Solanum melongena* L.): the representative solanum species indigenous to the old world. DNA Res 21: 649–660.

Hoff KL et al (2016) BRAKER1: Unsupervised RNA-Seq-based genome annotation with GeneMark-ET and AUGUSTUS. Bioinformatics 32: 767–769.

Huang S et al (2013) Draft genome of the kiwifruit *Actinidia chinensis*. Nat Comm 4: 2640.

Iorizzo M et al (2016) A high-quality carrot genome assembly provides new insights into carotenoid accumulation and asterid genome evolution. Nat Genet 48: 657–666.

Jaillon O et al (2009) The grapevine genome sequence suggests ancestral hexaploidization in major angiosperm phyla. Nature 449: 463–467.

Kazama Y et al (2016) A new physical mapping approach refines the sex-determining gene positions on the *Silene latifolia* Y-chromosome. Sci Rep 6: 18917.

Komatsuda T et al (2007) Six-rowed barley originated from a mutation in a homeodomain-leucine zipper I-class homeobox gene. Proc Nat Acad Sci USA 104: 1424–1429.

Krishnakumar V et al (2015) Araport: the Arabidopsis information portal. Nucl Acids Res 43: D1003–1009.

Khan, A., Fornes, O., Stigliani, A., Gheorghe, M., Castro-Mondragon, J.A., van der Lee, R., Bessy, A., Chèneby, J., Kulkarni, S.R., Tan, G., Baranasic, D., Arenillas, D.J., Sandelin, A., Vandepoele, K., Lenhard, B., Ballester, B., Wasserman, W.W., Parcy, F. and Mathelier, A. (2018) JASPAR 2018: update of the open-access database of transcription factor binding profiles and its web framework. Nucleic Acids Res.: D260–D266. https://doi.org/10.1093/nar/gkx1126

Krzywinski M et al (2009) Circos: an information aesthetic for comparative genomics. Genome Res 19: 1639–1645.

Langmead B, Salzberg SL (2012) Fast gapped-read alignment with Bowtie 2. Nat Methods 9: 357–359.

Li H et al (2009) The Sequence Alignment/Map format and SAMtools. Bioinformatics 25: 2078–2079.

Librado P, Rozas J (2009) DnaSP v5: a software for comprehensive analysis of DNA polymorphism data. Bioinformatics 25: 1451–1452.

Liu Z et al (2004) A primitive Y chromosome in papaya marks incipient sex chromosome evolution. Nature 427: 348–352.

Lomsadze A et al (2011) Improved prokaryotic gene prediction yields insights into transcription and translation mechanisms on whole genome scale. bioRxiv doi: https://doi.org/10.1101/193490.

Ming R, Bendahmane A, Renner SS (2011) Sex chromosomes in land plants. Annu Rev Plant Biol 62: 485–514.

Nettancourt DD (2001) Incompatibility and incongruity in wild and cultivated plants. Berlin: Springer-Verlag.

Olsen JL et al (2016) The genome of the seagrass *Zostera marina* reveals angiosperm adaptation to the sea. Nature 530: 331–335.

O’Malley, R.C., Huang, S.C., Song, L., Lewsey, M.G., Bartlett, A., Nery, J.R., Galli, M., Gallavotti, A. and Ecker, J.R. (2016) Cistrome and epicistrome features shape the regulatory DNA landscape. Cell, 165, 1280–1292.

Peterson BK et al (2012) Double digest RADseq: an inexpensive method for de novo SNP discovery and genotyping in model and non-model species. PLoS ONE 7: e37135.

Petersen SV, Dutton A, Lohmann KC (2016) End-Cretaceous extinction in Antarctica linked to both Deccan volcanism and meteorite impact via climate change. Nat Comm 7: 12079.

Price AL, Jones NC, Pevzner PA (2005) De novo identification of repeat families in large genomes. Bioinformatics 21: Suppl 1 i351–358.

Renner SS, Ricklefs RE (1995) Dioecy and its correlates in the flowering plants. Amer J Bot 82: 596–606.

Renner SS (2014) The relative and absolute frequencies of angiosperm sexual systems: dioecy, monoecy, gynodioecy, and an up-dated online database. Amer J Bot 101: 1588–1596.

Reyes-Chin-Wo S et al (2017) Genome assembly with in vitro proximity ligation data and whole-genome triplication in lettuce. Nat Comm 8: 14953.

Roulin A et al (2013) The fate of duplicated genes in a polyploid plant genome. Plant J 73: 143–153.

Sakuma S et al (2013) Divergence of expression pattern contributed to neofunctionalization of duplicated HD-Zip I transcription factor in barley. New Phytol 197: 939–948.

Sakuma S, Salomon B, Komatsuda T (2011) The domestication syndrome genes responsible for the major changes in plant form in the Triticeae crops. Plant Cell Physiol 52: 738–749.

Schmieder R, Edwards R (2011) Quality control and preprocessing of metagenomic datasets, Bioinformatics 27: 863–864.

Shi T, Huang H, Barker MS (2010) Ancient genome duplications during the evolution of kiwifruit (*Actinidia*) and related Ericales. Annals Bot 106: 497–504.

Shirasawa K, Hirakawa H, Isobe S (2016) Analytical workflow of double-digest restriction site-associated DNA sequencing based on empirical and *in silico* optimization in tomato. DNA Res 23: 145–153.

Shirasawa et al (2017) The genome sequence of sweet cherry (*Prunus avium*) for use in genomics-assisted breeding. DNA Res 24: 499–508.

Simão FA et al (2015) BUSCO: assessing genome assembly and annotation completeness with single-copy orthologs. Bioinformatics 31: 3210–3212.

Stanke M, Waack S (2003) Gene prediction with a Hidden-Markov model and a new intron submodel. Bioinformatics 19: 215–225.

Tamura K et al (2013) MEGA6: Molecular Evolutionary Genetics Analysis Version 6.0. Mol Biol Evol 30: 2725–2729.

Tamura M, Tao R, Yonemori K, Ustunomiya N, Sugiura A (1998) Ploidy level and genome size of several *Diospyros* species. J Jpn Soc Hortic Sci 67: 306–312.

The Tomato Genome Consortium (2012) The tomato genome sequence provides insights into fleshy fruit evolution. Nature 485: 635–641.

Trapnell C, Pachter L, Salzberg SL (2009) TopHat: discovering splice junctions with RNA-Seq. Bioinformatics 25: 1105–1111.

Tuskan GA et al (2006) The genome of black cottonwood, *Populus trichocarpa* (Torr. & Gray). Science 313: 1596–1604.

Van de Peer Y,Mizrachi E, Marchal K (2017) The evolutionary significance of polyploidy. Nat Rev Genet 18: 411–424.

Vanneste K, Baele G, Maere S, Van De Peer Y (2014) Analysis of 41 plant genomes supports a wave of successful genome duplications in association with the Cretaceous– Paleogene boundary. Genome Res 24: 1334–1347.

Wang J et al (2012) Sequencing papaya X and Yh chromosomes reveals molecular basis of incipient sex chromosome evolution. Proc Nat Acad Sci USA 109: 13710–13715.

Wilf P, Labandeira CC, Johnson KR, Ellis B (2006) Decoupled plant and insect diversity after the end-Cretaceous extinction. Science 313: 1112–1115.

Whipple CJ et al (2011) *grassy tillers1* promotes apical dominance in maize and responds to shade signals in the grasses. Proc Nat Acad Sci USA 108: E506–E512.

Yang HW, Akagi T, Kawakatsu T, Tao R (2019) Gene networks orchestrated by factor mechanism underlying sex determination in persimmon. Plant J 98: 97–111.

Yang Z (1997) PAML: a program package for phylogenetic analysis by maximum likelihood. Computer Appl Biosci 13: 555–556.

