## Supplemental Figures and Tables for "The persimmon genome reveals clues to the evolution of a lineage-specific sex determination system in plants"

### Supplementary Information

**Supplemental Figure 1: kmer distribution to estimate genome size and the degree of heterozygosity**

The distribution of distinct *k*-mers (k = 17) from the Illumina short reads showed two peaks at multiplicities of 32 and 54. The low and high peaks represent heterozygous and homozygous sequences, respectively. We estimated the genome size to be 877.7 Mb from the higher peak. This estimation almost agreed with the value measured by flow cytometry, 907 Mb, which was calculated from the nuclear DNA content in *D. lotus* of 1.85 pg/2 C (Tamura et al., 1998) and an assumption that 1 pg of DNA is equivalent to 980 Mb (Bennett et al., 2000).

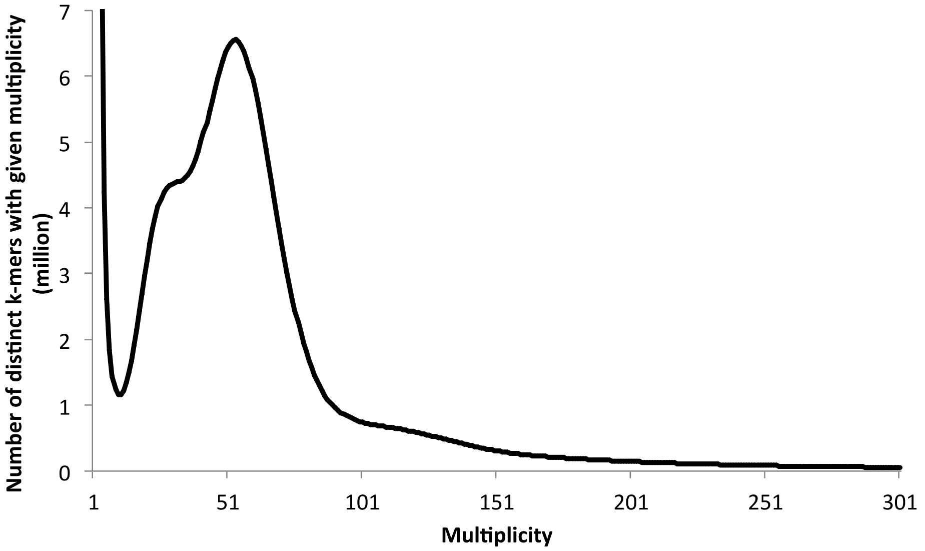

**Supplemental Figure 2: Genetic anchoring data (Dlo01-15).**

Genetic linkage map (left bars) and physical map of Dlo_r1.0 pseudomolecule sequences (right bars). Colors in the genetic map represent density of SNPs per 5 cM, while black, white and gray bars in the physical map indicate the positions of the forward, invert, and unknown directional scaffold sequences integrated in the pseudomolecule, respectively.

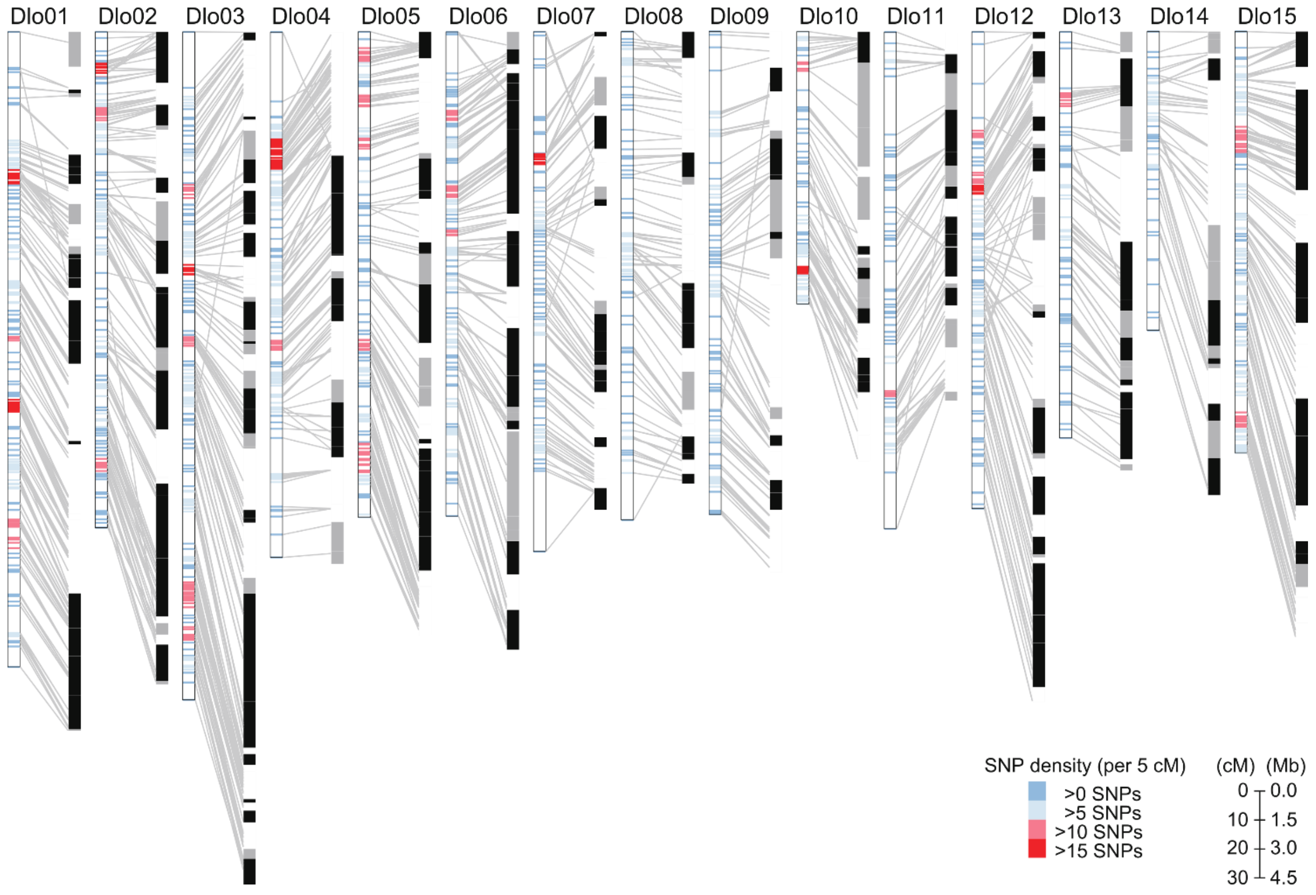

**Supplemental Figure 3: Conservation of gene and repetitive sequences across representative plant species**

**a,** Amino acid sequences were compared among genes from *D. lotus* (40,532 genes; DLO_r1.1 primary), *A. chinensis* (39,040 genes (Huang et al., 2013)), *V. vinifera* (29,927 genes (IGGP 12x.31) (Jaillon et al., 2009)), *S. lycopersicum* (34,789 genes (ITAG 3.10) (The Tomato Genome Consortium, 2012)), and *A. thaliana* (27,655 genes (Araport11) (Cheng et al., 2017)) using OrthoMCL v2.0.9 (Li et al., 2003) with default parameters. The numbers of clusters were shown in the intersections of the Venn diagram. **b,** Repetitive sequences were identified by RepeatMasker v4.0.6 (http://www.repeatmasker.org) using Repbase v406 (http://www.girinst.org/repbase/) and RepeatScout v1.0.5 for the genome sequences of *D. lotus* (8,974 sequences; DLO_r1.0), *A. chinensis* (30 pseudomolecules (Huang et al., 2013)), *S. lycopersicum* (13 pseudomolecules (SL3.0) (The Tomato Genome Consortium, 2012)), *Lactuca sativa* (lettuce; 9 pseudomolecules (V8) (Reyes-Chin-Wo et al., 2017)), *V. vinifera* (33 chromosomes (IGGP 12x.31) (Jaillon et al., 2009)), *Prunus persica* (peach; 8 pseudomolecules (v2.0.a1) (Verde et al., 2013), *Carica papaya* (papaya; 5,901 scaffold sequences (ASGPBv0.4) (Ming et al., 2008)), and *A. thaliana* (5 chromosomes, chloroplast and mitochondria genomes (TAIR10)). The percentage of repetitive sequences against the total length of the genome sequence were calculated for each of the result of RepeatMasker and RepeatScout and compared among the plant species.

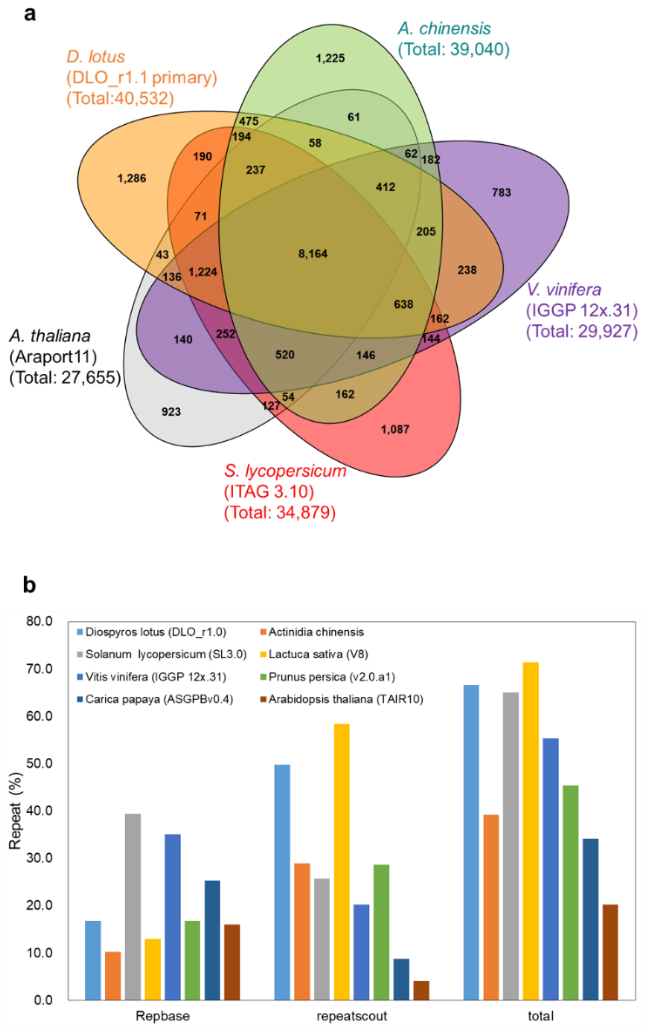

**Supplemental Figure 4: Gene duplication patterns following the *Dd-α* event.**

**a,** Heat map for the numbers of genes derived from *Dd-α*, shared between two chromosomes. For instance, Dlo01 shared many paralogs with Dlo02 and Dlo12, while Dlo02 and Dlo12 shared few paralogs with each other. The patterns of such affinities between the chromosomes suggests a paleotetraploidization event. **b,** Syntenic relationship betweem the putatively *Dd-α*-derived paralogous genes within the *Diospyros* genome. **c**, Synteny dot plot in a putative paleoduplicated region, on chromosomes 1 and 2. The red dotted boxes indicate long segmental syntenic blocks (>3Mb).

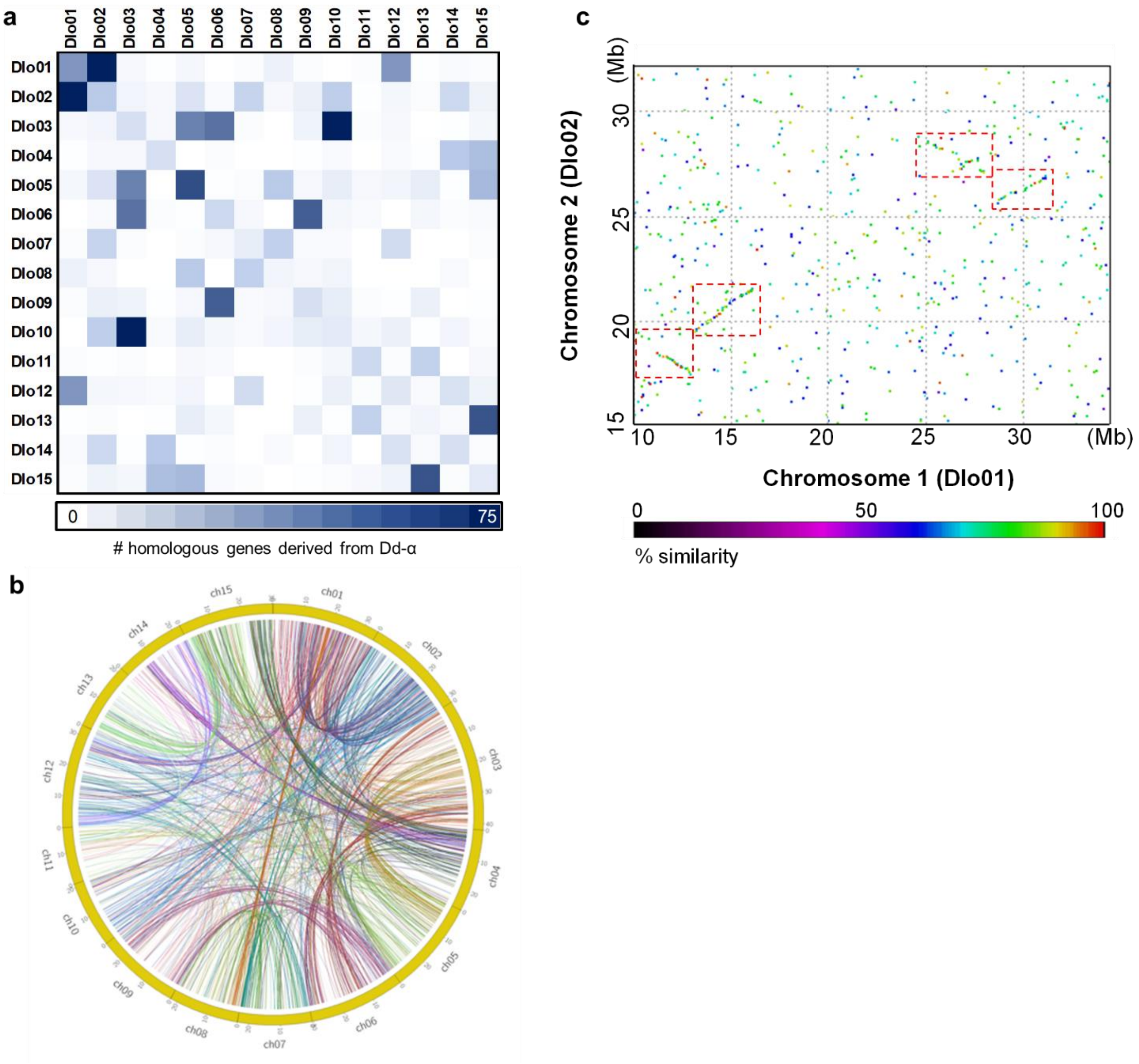

**Supplemental Figure 5: Genome-wide synteny between *Diospyros* and *Actinidia***

Dot plots of the syntenic genomic regions between *Diospyros* and *Actinidia*. As represented in the orange box, a single genomic segment from *Actinidia* corresponds to two syntenic *Diospyros* genome regions which are derived from the *Dd-α*. In the orange box, the middle regions of Dlo01 and Dlo02, and Dlo03 and Dlo06 are duplicated regions via *Dd-α* (see Figure 1g and Supplemental Figure 6). On the other hand, as represented in the green box, a genomic segment from *Diospyros* corresponds to at maximum four syntenic *Actinidia* genome regions which are derived from the double *Actinidia*-specific paleoduplication events (Ad-α and Ad-β) (Huang et al., 2013). These results indicate that, in the evolution of the order Ericales, *Dd-α* and Ad-α/β occurred independently in the *Diospyros* and *Actinidia* ancestral genomes, respectively.

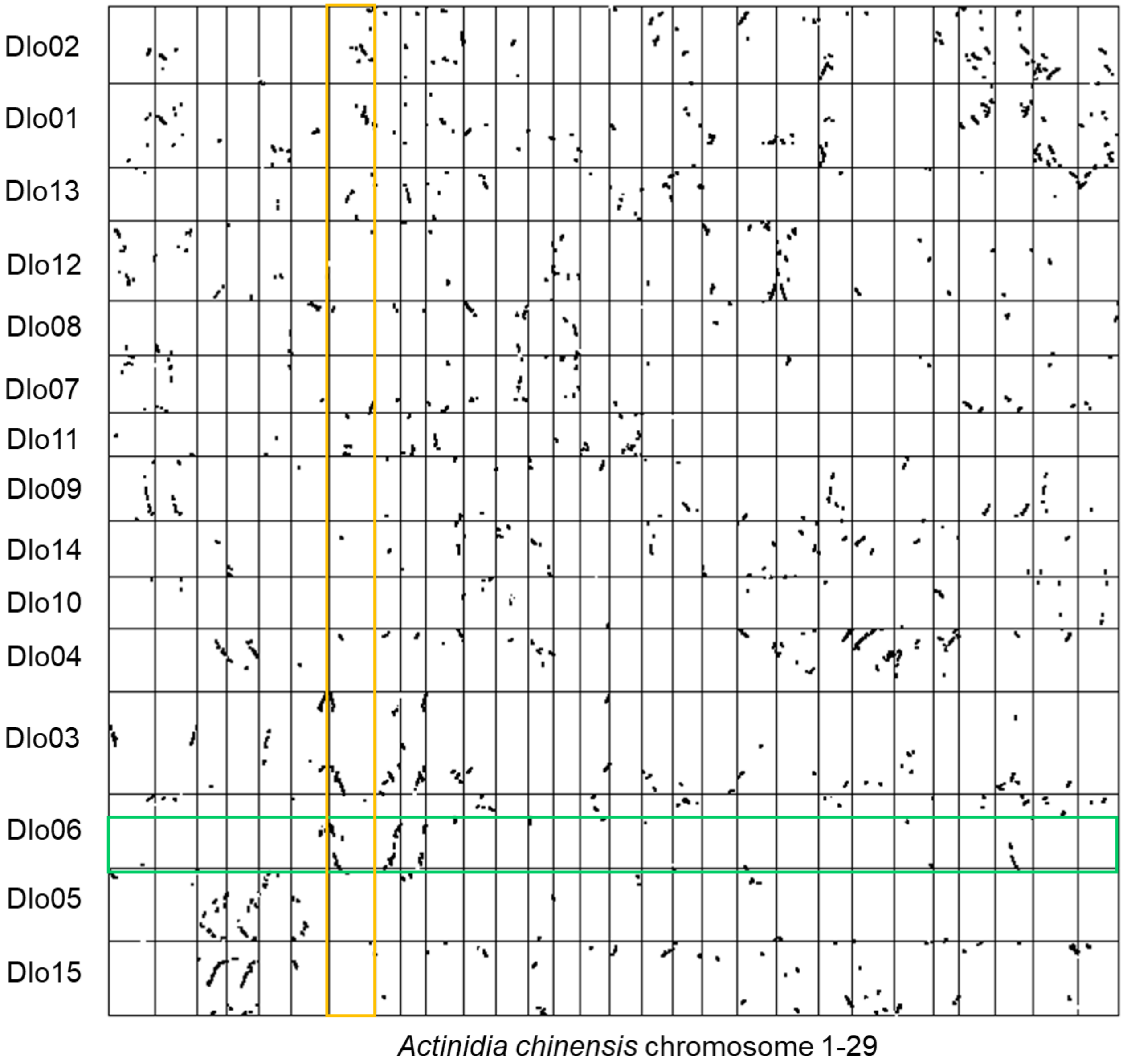

**Supplemental Figure 6: Phylogenetic relationship of wide HD-ZIP1 family in angiosperm.**

Phylogeny of the HD-Zip1 type homeodomain genes in representative angiosperm genomes (*Solanum lycopersicum, Oryza sativa, Zea mays*, and *Arabidopsis thaliana*) and *D. lotus* genome. The 175 HD-ZIP1 genes constituted at least 8 major clades (Clade I-VIII), which included at least one homologs from all 5 species used in this study (see Materials and Methods), although the root branches of the clade VI and VII were not statistically significant (32/100 and 28/100, respectively). MeGI, SiMeGI and Vrs1 were nested to the clade IV, given in light green.

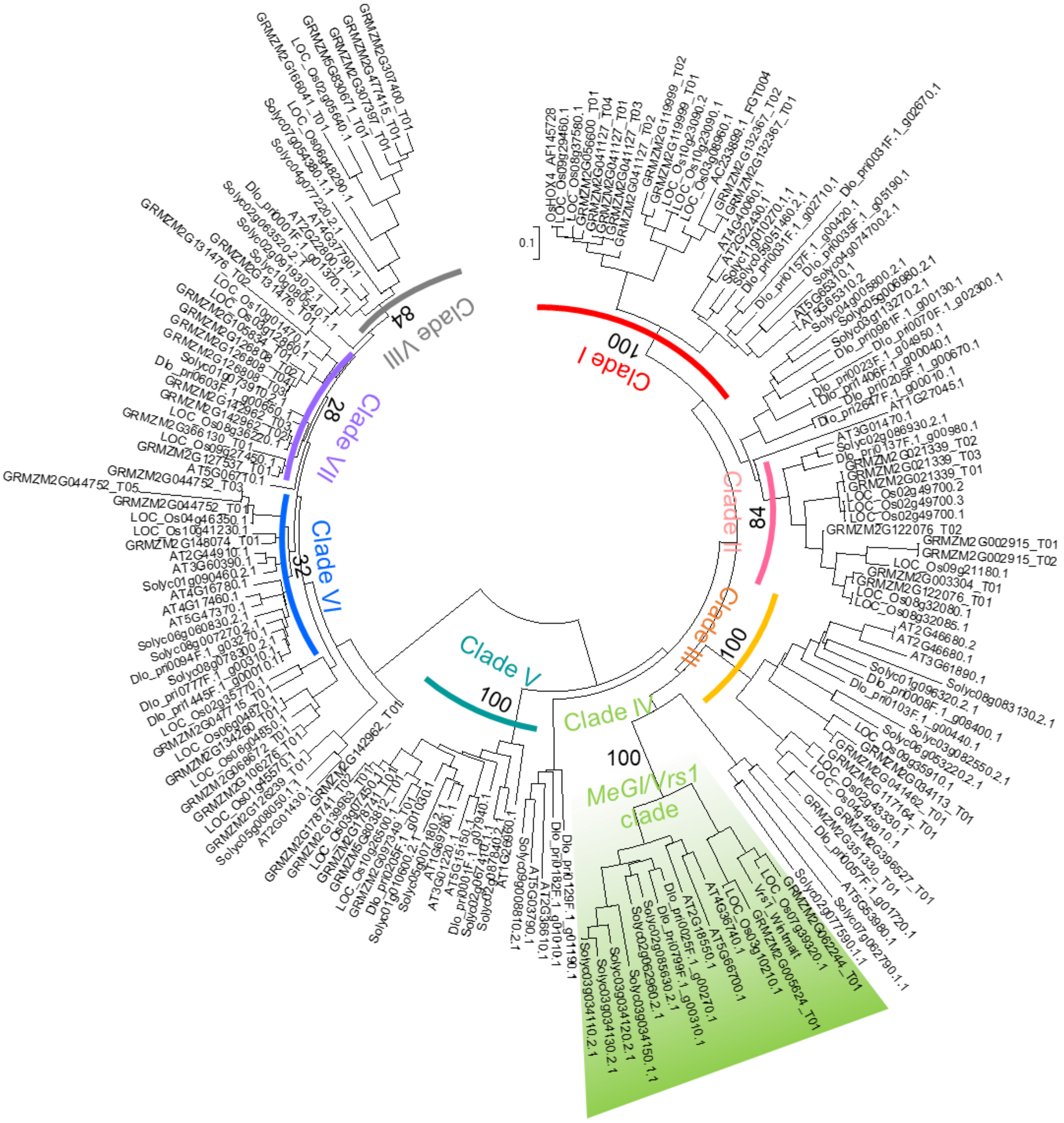

**Supplemental Figure 7: Overexpression of *MeGI* and *SiMeGI* under the control of CaMV35S promoter in *N. tabacum***

**a**-**c**, 1-week old transgenic lines. The *MeGI*-induced lines (**a**) frequently showed clear irregularities in development, in comparison to the *SiMeGI* lines (**b**) or empty cassette-induced lines (**c**). **d**, Comparison of 4-weeks old transgenic plants. The *MeGI*-induced lines (center) uniformly showed more severe growth inhibition, than the *SiMeGI*-induced lines (left). **e**, close-up picture of the *MeGI*-induced line corresponding to the individual marked with an asterisk in the panel (**d**). The leaves showed irregular shapes with significantly less veins. **f**, comparison of the appearance of 15-weeks old transgenic lines. The *MeGI*-induced line (left) exhibited dwarfism, but the total number of leaves were comparable to the control plants (right), while the internode lengths were shorter than the control, as shown in the panel "**g**". **h**, Differentiation of the leave shapes and structures in the control (left) and the *MeGI*-induced line (right). The *MeGI*-induced lines produced narrow and serrated leaves. Bars indicate 10mm for a-c, and e; 50mm for d, f, g, and h.

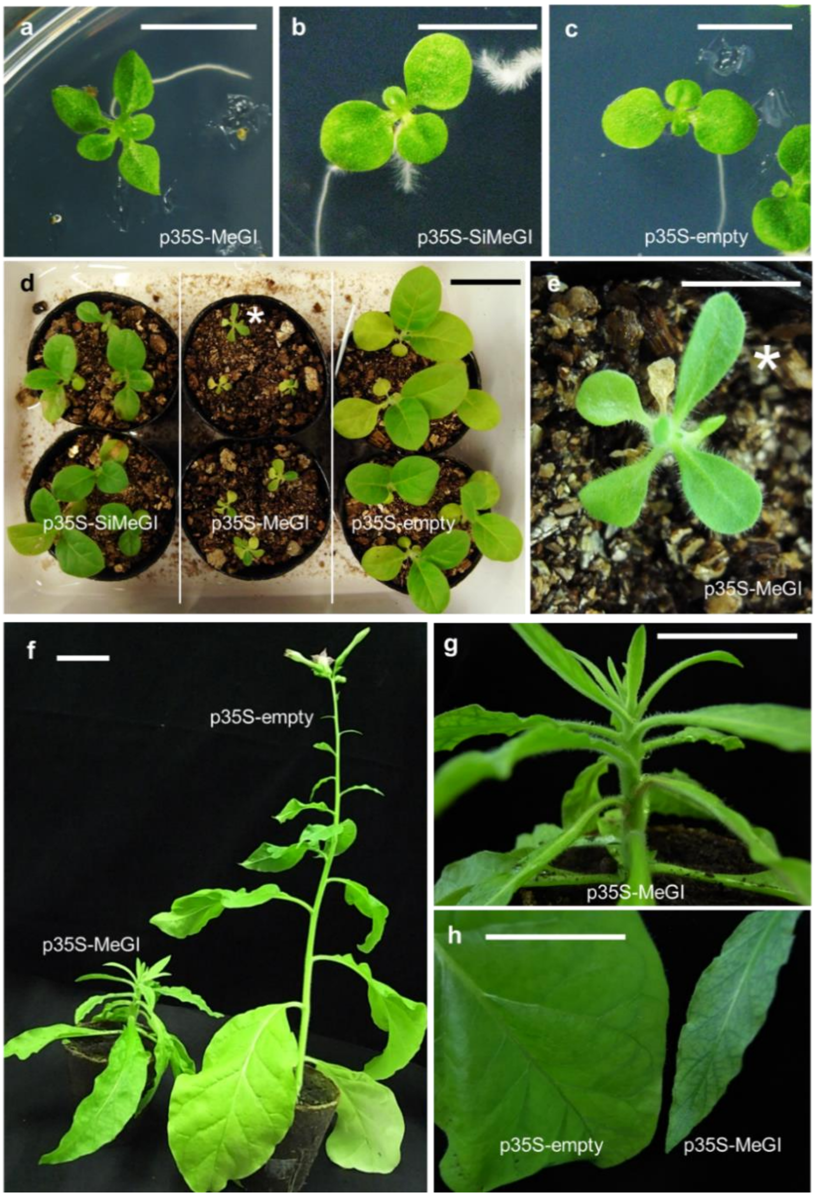

**Supplemental Figure 8: Overexpression of *MeGI* and *SiMeGI* under the control of CaMV35S promoter in *A. thaliana.***

**a**, Dissection of the control Arabidopsis plant transformed with an empty cassette. an: anther, pe: petal, sg: stigma. **b-e**, p35S-*MeGI* transgenic lines. Dissected flowers show rudimental anthers (ra) (**b-c**). Approximately half of the transgenic plants are semi-dwarf (semi-dwf) (**d**) or complete dwarf (**e**). They also frequently showed leaf serration, which is consistent with our previous analysis of the p35S-*MeGI* induced Arabidopsis plants (Akagi et al., 2014). **f-i**, p35S-*SiMeGI* transgenic lines. The transgenic plants occasionally showed rudimental anthers similar to the *MeGI*-induced lines (**f-g**). A part of the *SiMeGI*-induced lines showed semi-dwarfism (**h**), but full dwarfism was never observed in the 63 transgenic lines. Over 95% of the *SiMeGI*-induced lines were hermaphroditic, where the numbers of stamens are properly maintained (**i**), in contrast to the *MeGI*-induced lines (Akagi et al., 2014). Bars indicate 1mm for a, b, f, and i; 0.1mm for c and g; 10mm for d-e and h. j, Distribution of the number of female (fe) and hermaphrodite (herm) individuals in the p35-*MeGI* (green), p35S-*SiMeGI* (yellow), and p35S-empty (cont; gray) transgenic lines. k, Distribution of the number of complete dwarf (dwf), semi-dwarf (semi-dwf) and normal individuals in the p35-*MeGI*, p35S-*SiMeGI*, and p35S-empty (cont) transgenic lines.

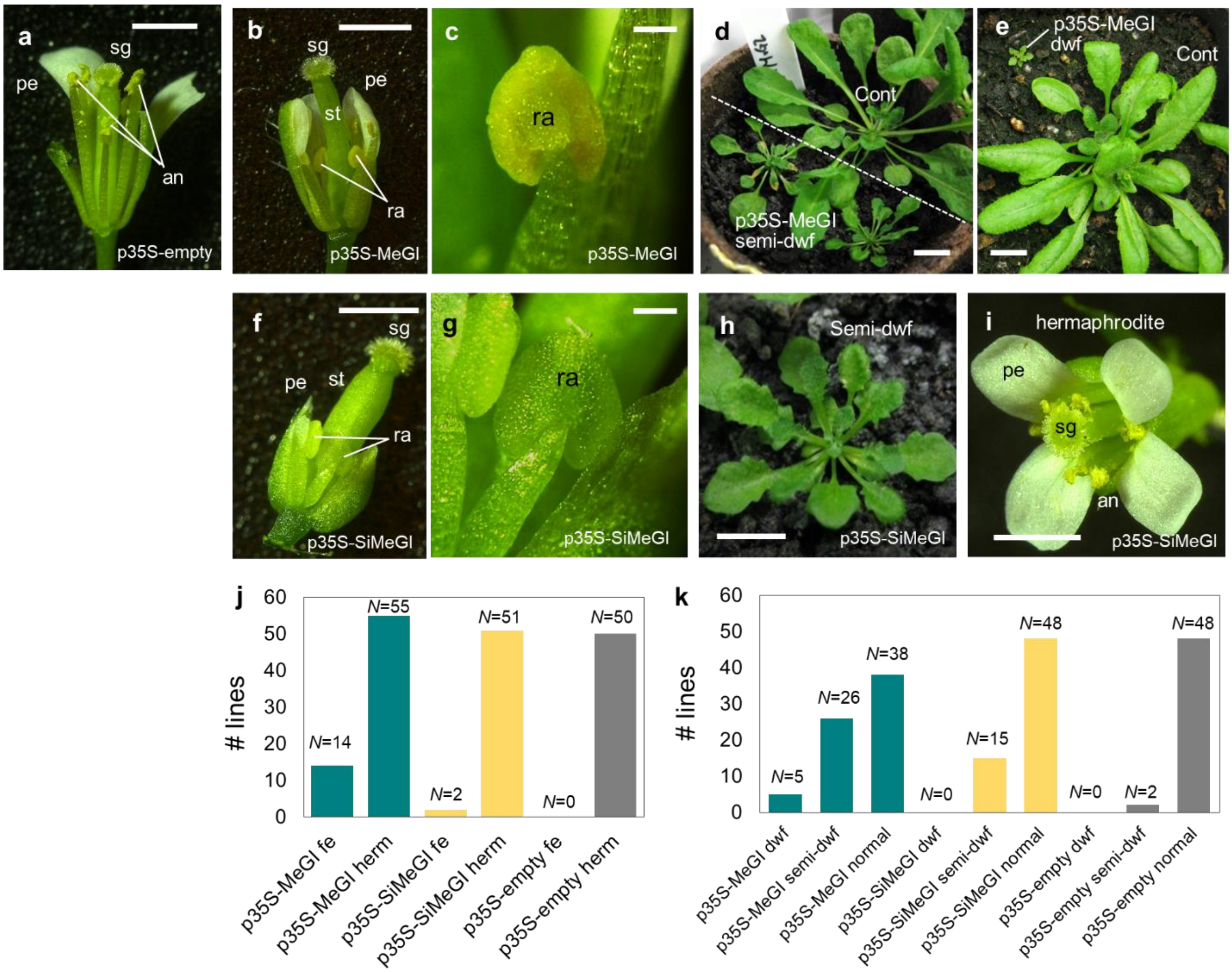

**Supplemental Figure 9: *in situ* RNA hybridization**

RNA *in situ* hybridization in developing buds and flower primordia, using *MeGI* (**a-c**) and *SiMeGI* (**d-f**) sequences as probes. In the cross section of the developing buds (**a** and **d**), the *MeGI* signal is strong in flower buds only (fb) (**a**), while *SiMeGI* showed significant signal in the pith (Pi) and young leaves (ly), as well as in flower buds (**d**). This is consistent with our expression analyses using laser capture micro-dissected (LCM) samples (Supplemental Figure 11). In the longitudinal sections of the developing buds (**b** and **e**), both *MeGI* and *SiMeGI* signals are confined to the meristematic region, especially in the shoot apical meristems (sam). At a later developing stage (**c** and **f**), flower primordia (fp) and bract (br) showed substantial signals of both *MeGI* (**c**) and *SiMeGI* (**f**). Bars indicate 50μm.

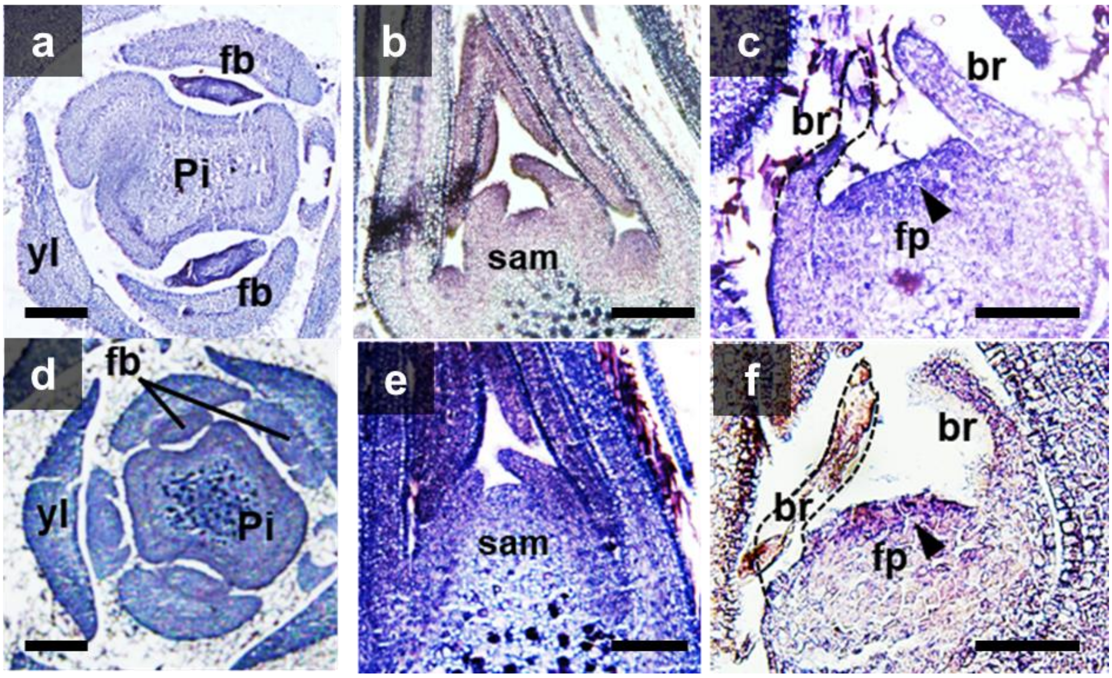

**Supplemental Figure 10: Expression analysis of *MeGI/SiMeGI* in laser capture microdissection (LCM)**

Longitudinal (**a**) and cross (**b**) sections of buds from *D. lotus*, Kunsenshi-male, and the target of the LCM. We targeted flower buds (red), young leaf or leaf buds (blue), and pith or cambium (green). **c**, the section after laser captions. **d**, qRT-PCR analysis to detect relative expression of the *MeGI* and *SiMeGI* among the organs, at early developmental stages (Jun-Jul) when the flower primordia form. Consistent with the results of the in situ hybridization (Supplemental Figure 10), *MeGI* expression was much stronger in flower buds than in pith or young leaves, while the difference in expression levels between the three organs was less drastic for *SiMeGI*. For both graphs, the expression level in flower buds was defined as “1”. **e**, comparison of the expression level of *MeGI* and *SiMeGI* in the developing flower buds. Illumina mRNA-Seq analysis was conducted on the LCM samples to detect RPKM values of *MeGI* and *SiMeGI*. *SiMeGI* was expressed higher than or comparative to the *MeGI,* in the developing flower buds. Notwithstanding, the reduction in *MeGI* expression in this stage affect the flower sexuality and the inflorescent structure (Akagi et al., 2014). **f**, relative expression of the *MeGI* and *SiMeGI* in different organs, during dormancy stage (Dec) when the development of flower primordia halt. Flower buds showed no significant expression of either *MeGI* or *SiMeGI*.

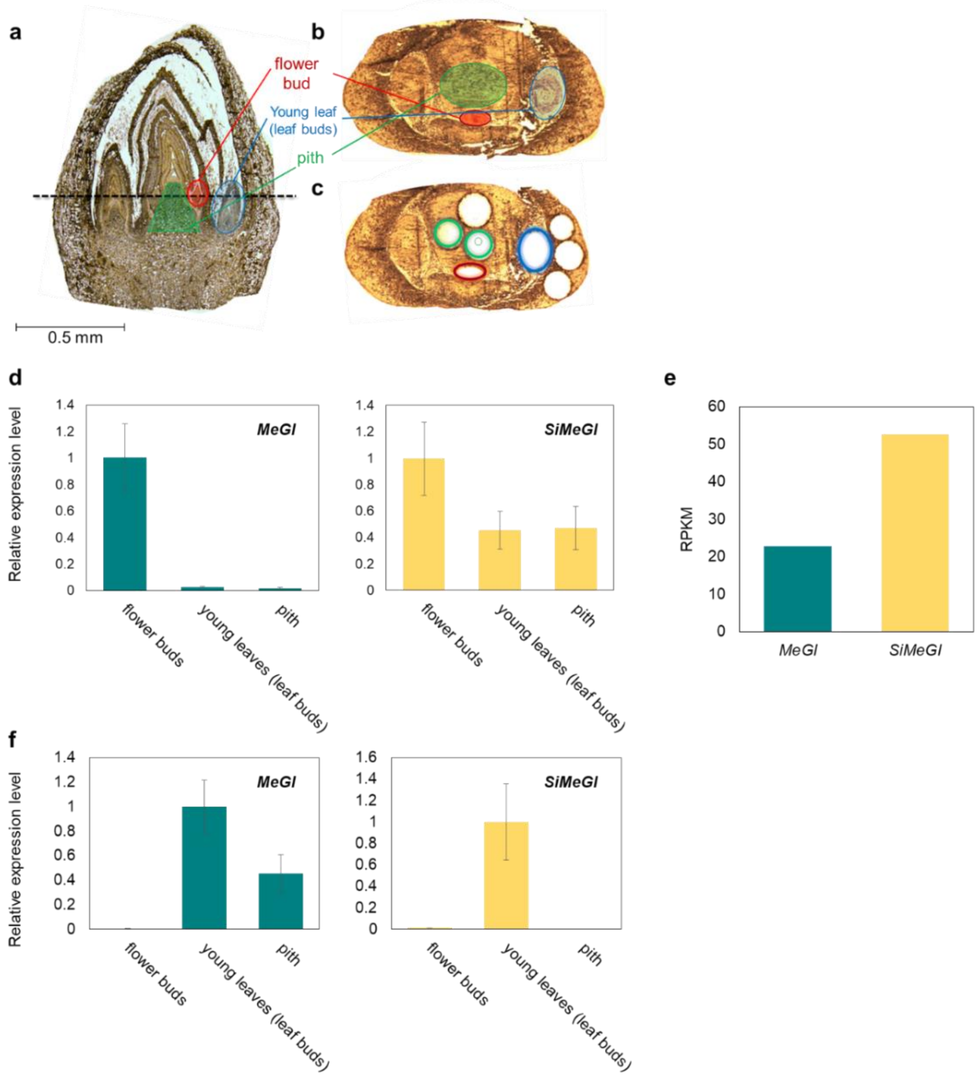

### S1 Table: Summary statistics for the initial genome assembly of *D. lotus* cv. Kunsenshi-male

| | | *Diospyros lotus* (DLO_r1.0) | *Diospyros lotus* (DLO_r1.0p) | *Diospyros lotus* (DLO_r1.0a) |
|---|---|---|---|---|
| Estimated genome size | | - | 877.7 Mb | - |
| Total | Number of sequences | 8,974 | 3,073 | 5,901 |
| | Total length (bp) | 945,625,305 | 746,093,016 | 199,532,289 |
| | Average length (bp) | 105,374 | 242,790 | 33,813 |
| | Max length (bp) | 7,107,382 | 7,107,382 | 505,402 |
| | Min length (bp) | 2,483 | 2,483 | 2,685 |
| | N50 length (bp) | 653,828 | 1,060,344 | 43,117 |
| | A | 300,991,439 | 237,381,743 | 63,609,696 |
| | T | 300,899,593 | 237,282,679 | 63,616,914 |
| | G | 171,753,801 | 135,616,131 | 36,137,670 |
| | C | 171,980,472 | 135,812,463 | 36,168,009 |
| | n | 0 | 0 | 0 |
| | N | 0 | 0 | 0 |
| | others | 0 | 0 | 0 |
| | Total(ACGT) | 945,625,305 | 746,093,016 | 199,532,289 |
| | GC%(ACGT) | 36.3 | 36 | 36 |
| | N% | 0.0 | 0 | 0 |
| | (Estimated size-Total(ATGC)/Estimated size | - | 15.0 | - |
| ≥100 bp | Number of sequences | 8,974 | 3,073 | 5,901 |
| | Total length (bp) | 945,625,305 | 746,093,016 | 199,532,289 |
| | Average length (bp) | 105,374 | 242,790 | 33,813 |
| ≥200 bp | Number of sequences | 8,974 | 3,073 | 5,901 |
| | Total length (bp) | 945,625,305 | 746,093,016 | 199,532,289 |
| | Average length (bp) | 105,374 | 242,790 | 33,813 |
| ≥300 bp | Number of sequences | 8,974 | 3,073 | 5,901 |
| | Total length (bp) | 945,625,305 | 746,093,016 | 199,532,289 |
| | Average length (bp) | 105,374 | 242,790 | 33,813 |
| ≥400 bp | Number of sequences | 8,974 | 3,073 | 5,901 |
| | Total length (bp) | 945,625,305 | 746,093,016 | 199,532,289 |
| | Average length (bp) | 105,374 | 242,790 | 33,813 |
| ≥200 bp | Number of sequences | 8,974 | 3,073 | 5,901 |
| | Total length (bp) | 945,625,305 | 746,093,016 | 199,532,289 |
| | Average length (bp) | 105,374 | 242,790 | 33,813 |
| ≥1 kbp | Number of sequences | 8,974 | 3,073 | 5,901 |
| | Total length (bp) | 945,625,305 | 746,093,016 | 199,532,289 |
| | Average length (bp) | 105,374 | 242,790 | 33,813 |
| ≥2 kbp | Number of sequences | 8,974 | 3,073 | 5,901 |
| | Total length (bp) | 945,625,305 | 746,093,016 | 199,532,289 |
| | Average length (bp) | 105,374 | 242,790 | 33,813 |
| ≥3 kbp | Number of sequences | 8,964 | 3,070 | 5,894 |
| | Total length (bp) | 945,597,159 | 746,084,791 | 199,512,368 |
| | Average length (bp) | 105,488 | 243,024 | 33,850 |
| ≥4 kbp | Number of sequences | 8,883 | 3,042 | 5,841 |
| | Total length (bp) | 945,311,437 | 745,985,214 | 199,326,223 |
| | Average length (bp) | 106,418 | 245,229 | 34,125 |
| ≥5 kbp | Number of sequences | 8,805 | 3,021 | 5,784 |
| | Total length (bp) | 944,956,722 | 745,890,657 | 199,066,065 |
| | Average length (bp) | 107,320 | 246,902 | 34,417 |
| Estimated genome size | | - | 877,700,000 | - |

**S2 Table: Number of SNPs and length of genetic linkage maps in *D. lotus***

Genetic maps for the four parental lines of the two mapping populations (KK and VM), were built using the pseudo-test cross method. All linkage groups were anchored to the 15 chromosomes of the *D. lotus* draft genome assembly. The sex-determinant locus was mapped to linkage group 15, suggesting the Dlo15 is the sex chromosome. More detailed information about the maps and SNPs are available from the Persimmon Genome Database (http://persimmon.kazusa.or.jp)

| Chromsome name | Consensus map | | KK genetic map | | | | VM genetic map | | | |
|---|---|---|---|---|---|---|---|---|---|---|
| | | | Kunseishi-female | | Kunseishi-male | | Kunseishi-female | | Budogaki-male | |
| | No. of SNPs | Map length (cM) | No. of SNPs | Map length (cM) | No. of SNPs | Map length (cM) | No. of SNPs | Map length (cM) | No. of SNPs | Map length (cM) |
| Dlo01 | 435 | 223.8 | 140 | 218.4 | 204 | 208.5 | 154 | 176.0 | 144 | 110.6 |
| Dlo02 | 723 | 132.4 | 307 | 157.9 | 175 | 162.8 | 148 | 183.1 | 316 | 132.1 |
| Dlo03 | 675 | 63.9 | 236 | 134.4 | 222 | 219.4 | 238 | 177.4 | 311 | 166.8 |
| Dlo04 | 385 | 159.2 | 116 | 145.1 | 176 | 172.5 | 174 | 138.6 | 168 | 139.4 |
| Dlo05 | 384 | 136.4 | 106 | 103.7 | 180 | 159.5 | 191 | 191.5 | 133 | 136.4 |
| Dlo06 | 384 | 139.7 | 75 | 82.6 | 160 | 159.0 | 159 | 136.0 | 216 | 129.7 |
| Dlo07 | 262 | 139.6 | 97 | 122.3 | 124 | 170.7 | 81 | 97.7 | 94 | 114.2 |
| Dlo08 | 248 | 127.3 | 62 | 125.5 | 108 | 160.4 | 108 | 195.0 | 114 | 135.7 |
| Dlo09 | 279 | 99.6 | 58 | 105.9 | 103 | 158.5 | 115 | 110.1 | 155 | 115.2 |
| Dlo10 | 287 | 105.5 | 103 | 104.4 | 96 | 89.4 | 74 | 85.7 | 155 | 124.4 |
| Dlo11 | 395 | 147.1 | 150 | 147.1 | 96 | 163.1 | 35 | 85.6 | 213 | 160.7 |
| Dlo12 | 549 | 126.4 | 286 | 105.9 | 149 | 156.5 | 168 | 194.8 | 164 | 79.9 |
| Dlo13 | 187 | 49.5 | 36 | 139.2 | 84 | 133.4 | 53 | 77.2 | 109 | 99.4 |
| Dlo14 | 310 | 116.4 | 67 | 116.4 | 58 | 98.1 | 57 | 181.0 | 208 | 169.1 |
| Dlo15[a] | 456 | 146.5 | 148 | 121.7 | 185 | 138.4 | 202 | 166.2 | 189 | 129.0 |
| Total | 5,959 | 1,913.2 | 1,987 | 1,930.6 | 2,120 | 2,350.2 | 1,957 | 2,195.7 | 2,689 | 1,942.5 |

[a]Putative *D. lotus* sex chromosome, including the sex-determinant locus.

**S3 Table: Number of annotated SNPs and indels between the female and male lines of *D. lotus* 'Kunsenshi'**

SNPs and indels were identified from whole-genome resequencing analysis of female and male lines of *D. lotus*, and functionally annotated and classified into four categories predefined by SnpEff (Cingolani et al., 2012): high (e.g. nonsense mutations and frameshift mutations)-, moderate (e.g. missense mutations)-, modifier (e.g. intron and intergenic mutations)- and low-impact (e.g. synonymous mutations) mutations (see http://snpeff.sourceforge.net for details). Further details about these SNPs and Indels are available from the Persimmon Genome Database (http://persimmon.kazusa.or.jp).

| Annotation | Impact | #Variants | % |
|---|---|---|---|
| Stop gained | HIGH | 705 | >0.0% |
| Frameshift variant | HIGH | 641 | >0.0% |
| Stop lost & splice region variant | HIGH | 184 | >0.0% |
| Splice donor variant & intron variant | HIGH | 150 | >0.0% |
| Splice acceptor variant & intron variant | HIGH | 149 | >0.0% |
| Start lost | HIGH | 106 | >0.0% |
| Frameshift variant & splice region variant | HIGH | 33 | >0.0% |
| Frameshift variant & start lost | HIGH | 23 | >0.0% |
| Splice donor variant & splice region variant & intron variant | HIGH | 23 | >0.0% |
| Stop gained & inframe insertion | HIGH | 17 | >0.0% |
| Frameshift variant & stop lost & splice region variant | HIGH | 16 | >0.0% |
| Stop gained & splice region variant | HIGH | 16 | >0.0% |
| Frameshift variant & stop gained | HIGH | 11 | >0.0% |
| Stop gained & disruptive inframe insertion | HIGH | 8 | >0.0% |
| Splice acceptor variant & splice region variant & intron variant | HIGH | 3 | >0.0% |
| Stop lost & inframe deletion & splice region variant | HIGH | 2 | >0.0% |
| Frameshift variant & splice acceptor variant & splice region variant & intron variant | HIGH | 1 | >0.0% |
| Frameshift variant & splice donor variant & splice region variant & intron variant | HIGH | 1 | >0.0% |
| Start lost & disruptive inframe insertion | HIGH | 1 | >0.0% |
| Stop gained & disruptive inframe deletion | HIGH | 1 | >0.0% |
| Subtotal of high impact variants | | 2,091 | 0.1% |
| Missense variant | MODERATE | 27,843 | 1.6% |
| Inframe insertion | MODERATE | 404 | >0.0% |
| Missense variant & splice region variant | MODERATE | 377 | >0.0% |
| Inframe deletion | MODERATE | 199 | >0.0% |
| Disruptive inframe insertion | MODERATE | 191 | >0.0% |
| Disruptive inframe deletion | MODERATE | 114 | >0.0% |
| Inframe insertion & splice region variant | MODERATE | 7 | >0.0% |
| Disruptive inframe insertion & splice region variant | MODERATE | 5 | >0.0% |
| Disruptive inframe deletion & splice region variant | MODERATE | 1 | >0.0% |
| Subtotal of moderate impact variants | | 29,141 | 1.7% |
| Intergenic region | MODIFIER | 1,424,741 | 82.1% |
| Intron variant | MODIFIER | 223,388 | 12.9% |
| Intragenic variant | MODIFIER | 22,774 | 1.3% |
| Subtotal of modifier impact variants | | 1,670,903 | 96.3% |
| Synonymous variant | LOW | 24,127 | 1.4% |
| Splice region variant & intron variant | LOW | 2,928 | 0.2% |
| Splice region variant & synonymous variant | LOW | 363 | >0.0% |
| Splice region variant & stop retained variant | LOW | 64 | >0.0% |
| Initiator codon variant | LOW | 10 | >0.0% |
| Subtotal of low impact variants | | 27,492 | 1.6% |
| Unassigned | - | 5,656 | 0.3% |
| Total | | 1,735,283 | 100.0% |

**S4 Table: Comparison of the repeat sequences in representative eudicot genomes**

Repetitive sequences amounted for 630.2 Mb (66.6%) of the total length of the final genome assembly. Unique repeats were abundant in the *D. lotus* genomes, constituting 49.8% of all repeats. Of the known types of repeats, Class I LTR elements were observed most frequently (11.2%).

| | | | | | *Diospyros lotus* (DLO_r1.0) | | *Diospyros lotus* (DLO_r1.0p) | | *Diospyros lotus* (DLO_r1.0a) | | *Actinidia chinensis* | | *Solanum lycopersicum* (SL3.0) | |
|---|---|---|---|---|---|---|---|---|---|---|---|---|---|---|
| | | | | | Length occupied (bp) | % of whole genome | Length occupied (bp) | % of whole genome | Length occupied (bp) | % of whole genome | Length occupied (bp) | % of whole genome | Length occupied (bp) | % of whole genome |
| Known repeats | Interspersed repeats | Class I | SINEs | Total | 166,838 | 0.0 | 115,389.0 | 0.0 | 51,449.0 | 0.0 | 147,040 | 0.0 | 1,180,093 | 0.1 |
| | | | LINEs | Total | 7,806,418 | 0.8 | 6,073,244.0 | 0.8 | 1,733,174.0 | 0.9 | 1,968,880 | 0.3 | 10,156,031 | 1.2 |
| | | | LTR elements | Total | 106,069,966 | 11.2 | 83,669,799.0 | 11.2 | 22,400,167.0 | 11.2 | 37,945,569 | 6.0 | 268,611,657 | 32.4 |
| | | | | Copia | 57,552,037 | 6.1 | 45,352,029.0 | 6.1 | 12,200,008.0 | 6.1 | 20,917,900 | 3.3 | 43,976,597 | 5.3 |
| | | | | Gypsy | 46,703,277 | 4.9 | 36,947,330.0 | 5.0 | 9,755,947.0 | 4.9 | 15,343,487 | 2.4 | 219,399,447 | 26.5 |
| | | Class II | DNA elements | | 11,794,037 | 1.2 | 9,031,535.0 | 1.2 | 2,762,502.0 | 1.4 | 5,578,239 | 0.9 | 26,792,666 | 3.2 |
| | | Unclassified | | | 614 | 0.0 | 614.0 | 0.0 | 0.0 | 0.0 | 119 | 0.0 | 2,123,585 | 0.3 |
| | Helitrons | | | | 1,001,867 | 0.1 | 768,779.0 | 0.1 | 233,088.0 | 0.1 | 379,702 | 0.1 | 538,934 | 0.1 |
| | Low complexity | | | | 7,064,999 | 0.7 | 5,553,583.0 | 0.7 | 1,511,416.0 | 0.8 | 4,158,182 | 0.7 | 2,158,535 | 0.3 |
| | Simple repeat | | | | 22,555,879 | 2.4 | 17,650,019.0 | 2.4 | 4,905,860.0 | 2.5 | 12,809,197 | 2.0 | 9,387,824 | 1.1 |
| | Unknown | | | | 40,997 | 0.0 | 28,463.0 | 0.0 | 12,534.0 | 0.0 | 33,884 | 0.0 | 18,192 | 0.0 |
| | Subtotal | | | | 159,346,866 | 16.9 | 125,065,495.0 | 16.8 | 34,281,371.0 | 17.2 | 64,988,969 | 10.3 | 326,352,654 | 39.4 |
| Unique repeats | Unknown | | | | 470,242,683 | 49.7 | 371,663,642.0 | 49.8 | 98,579,041.0 | 49.4 | 182,375,737 | 28.9 | 212,578,211 | 25.7 |
| | Simple repeat | | | | 652,914 | 0.1 | 507,792.0 | 0.1 | 145,122.0 | 0.1 | 231,950 | 0.0 | 197,855 | 0.0 |
| | Subtotal | | | | 470,895,597 | 49.8 | 372,171,434.0 | 49.9 | 98,724,163.0 | 49.5 | 182,607,687 | 28.9 | 212,776,066 | 25.7 |
| | | | | | | | 497,236,929.0 | 66.6 | 133,005,534.0 | 66.7 | 247,596,656 | 39.2 | 539,128,720 | 65.1 |

| *Lactuca sativa* (V8) | | *Vitis vinifera* (IGGP 12x.31) | | *Prunus persica* (v2.0.a1) | | *Carica papaya* (ASGPBv0.4) | |
|---|---|---|---|---|---|---|---|
| Length occupied (bp) | % of whole genome | Length occupied (bp) | % of whole genome | Length occupied (bp) | % of whole genome | Length occupied (bp) | % of whole genome |
| 8,299 | 0.0 | 44,308 | 0.0 | 13,945 | 0.0 | 4,507 | 0.0 |
| 1,000,875 | 0.0 | 19,332,776 | 4.0 | 1,047,154 | 0.5 | 5,273,047 | 1.5 |
| 243,368,347 | 10.5 | 110,004,991 | 22.6 | 21,152,416 | 9.3 | 72,920,228 | 21.3 |
| 105,761,119 | 4.6 | 40,155,666 | 8.3 | 9,297,257 | 4.1 | 13,133,896 | 3.8 |
| 132,932,224 | 5.8 | 66,806,791 | 13.7 | 10,567,311 | 4.6 | 59,336,577 | 17.3 |
| 7,603,516 | 0.3 | 27,130,274 | 5.6 | 9,021,688 | 4.0 | 1,355,568 | 0.4 |
| 725 | 0.0 | 97 | 0.0 | 113 | 0.0 | 0 | 0.0 |
| 1,362,310 | 0.1 | 621,626 | 0.1 | 508,703 | 0.2 | 103,304 | 0.0 |
| 6,644,298 | 0.3 | 2,548,079 | 0.5 | 1,114,697 | 0.5 | 1,305,362 | 0.4 |
| 34,623,944 | 1.5 | 7,999,159 | 1.6 | 3,966,247 | 1.7 | 5,147,395 | 1.5 |
| 18,104 | 0.0 | 9,894 | 0.0 | 42,885 | 0.0 | 15,187 | 0.0 |
| 301,067,075 | 13.0 | 170,906,209 | 35.1 | 38,190,482 | 16.8 | 86,709,638 | 25.3 |
| 1,349,479,454 | 58.4 | 97,889,702 | 20.1 | 65,064,360 | 28.6 | 30,037,776 | 8.8 |
| 353,594 | 0.0 | 307,310 | 0.1 | 31,221 | 0.0 | 171,095 | 0.0 |
| 1,349,833,048 | 58.4 | 98,197,012 | 20.2 | 65,095,581 | 28.6 | 30,208,871 | 8.8 |

| *Arabidopsis thaliana* (TAIR10) | |
|---|---|
| Length occupied (bp) | % of whole genome |
| 123,259 | 0.1 |
| 1,336,827 | 1.1 |
| 8,154,990 | 6.8 |
| 1,818,720 | 1.5 |
| 6,224,830 | 5.2 |
| 5,585,341 | 4.7 |
| 0 | 0.0 |
| 2,173,124 | 1.8 |
| 406,058 | 0.3 |
| 1,314,108 | 1.1 |
| 6,400 | 0.0 |
| 19,220,552 | 16.1 |
| 4,935,265 | 4.1 |
| 10,315 | 0.0 |
| 4,945,580 | 4.1 |
| 24,166,132 | 20.2 |

### S5 Table: Phenotypic characterization of the p35S-*MeGI N. tabacum* transformed lines.

| T1 Line ID | introduced construct | feminization[a] | narrow leaves[b] | dwarfisms[c] | transgene expression in flowers |
|---|---|---|---|---|---|
| Nita-p35S-MeGI-1 | pGWB2-MeGI | no flowers | ++ | ++ | + |
| Nita-p35S-MeGI-2 | pGWB2-MeGI | — | — | — | + |
| Nita-p35S-MeGI-3 | pGWB2-MeGI | + | + | — | + |
| Nita-p35S-MeGI-4 | pGWB2-MeGI | + | + | + | + |
| Nita-p35S-MeGI-5 | pGWB2-MeGI | + | — | — | + |
| Nita-p35S-MeGI-6 | pGWB2-MeGI | + | + | — | + |
| Nita-p35S-MeGI-7 | pGWB2-MeGI | + | + | + | + |
| Nita-p35S-MeGI-8 | pGWB2-MeGI | — | — | — | + |
| Nita-p35S-MeGI-9 | pGWB2-MeGI | + | — | — | + |
| Nita-p35S-MeGI-10 | pGWB2-MeGI | — | — | — | — |
| Nita-p35S-MeGI-11 | pGWB2-MeGI | — | + | — | + |
| Nita-p35S-MeGI-12 | pGWB2-MeGI | no flowers | ++ | ++ | + |
| Nita-p35S-MeGI-13 | pGWB2-MeGI | — | + | — | + |
| Nita-p35S-MeGI-14 | pGWB2-MeGI | + | — | — | + |
| Nita-p35S-MeGI-15 | pGWB2-MeGI | + | — | — | + |
| Nita-p35S-MeGI-16 | pGWB2-MeGI | + | — | — | + |
| Nita-p35S-MeGI-17 | pGWB2-MeGI | + | + | + | + |
| Nita-p35S-MeGI-18 | pGWB2-MeGI | no flowers | ++ | ++ | + |
| Nita-p35S-MeGI-19 | pGWB2-MeGI | + | + | + | + |
| Nita-p35S-MeGI-20 | pGWB2-MeGI | — | + | + | + |
| Nita-p35S-MeGI-21 | pGWB2-MeGI | — | — | — | + |
| Nita-p35S-MeGI-22 | pGWB2-MeGI | + | ++ | + | + |
| Nita-p35S-MeGI-23 | pGWB2-MeGI | + | ++ | + | + |

[a] "+" indicates feminization.
[b] "+" indicates narrow leaves, as shown in Figure 3 and Figure S8.
[c] "+" and "++" indicate semi-dwarfing and dwarfing phenotypes, respectively, as shown in Figure 3.

### S6 Table: Phenotypic characterization of the p35S-*SiMeGI N. tabacum* transformed lines.

| T1 Line ID | introduced construct | feminization[a] | narrow leaves[b] | dwarfisms[c] | transgene expression in flowers |
|---|---|---|---|---|---|
| Nita-p35S-SiMeGI-1 | pGWB2-SiMeGI | — | — | + | + |
| Nita-p35S-SiMeGI-2 | pGWB2-SiMeGI | — | — | + | + |
| Nita-p35S-SiMeGI-3 | pGWB2-SiMeGI | — | — | — | — |
| Nita-p35S-SiMeGI-4 | pGWB2-SiMeGI | — | — | — | + |
| Nita-p35S-SiMeGI-5 | pGWB2-SiMeGI | — | — | + | + |
| Nita-p35S-SiMeGI-6 | pGWB2-SiMeGI | — | — | + | + |
| Nita-p35S-SiMeGI-7 | pGWB2-SiMeGI | — | — | — | + |
| Nita-p35S-SiMeGI-8 | pGWB2-SiMeGI | — | — | — | + |
| Nita-p35S-SiMeGI-9 | pGWB2-SiMeGI | — | — | — | + |
| Nita-p35S-SiMeGI-10 | pGWB2-SiMeGI | — | — | — | + |
| Nita-p35S-SiMeGI-11 | pGWB2-SiMeGI | — | — | — | + |
| Nita-p35S-SiMeGI-12 | pGWB2-SiMeGI | — | — | + | + |
| Nita-p35S-SiMeGI-13 | pGWB2-SiMeGI | — | — | — | + |
| Nita-p35S-SiMeGI-14 | pGWB2-SiMeGI | — | — | + | + |
| Nita-p35S-SiMeGI-15 | pGWB2-SiMeGI | — | — | — | + |
| Nita-p35S-SiMeGI-16 | pGWB2-SiMeGI | — | — | — | + |
| Nita-p35S-SiMeGI-17 | pGWB2-SiMeGI | — | — | — | — |
| Nita-p35S-SiMeGI-18 | pGWB2-SiMeGI | — | — | + | + |
| Nita-p35S-SiMeGI-19 | pGWB2-SiMeGI | — | — | + | + |
| Nita-p35S-SiMeGI-20 | pGWB2-SiMeGI | — | — | + | + |

[a] "+" indicates feminization.
[b] "+" indicates narrow leaves, as shown in Figure 3.
[c] "+" indicates semi-dwarf phenotype, as shown in Figure 3.

### S7 Table: Phenotypic characterization of the p35S-*MeGI A. thaliana* transformed lines.

| T1 Line ID | introduced construct | feminization[a] | dwarfism[b] | transgene expression in whole plant[c] |
|---|---|---|---|---|
| Arth-p35S-MeGI-1 | pGWB2-MeGI | + | ++ | ++ |
| Arth-p35S-MeGI-2 | pGWB2-MeGI | + | + | ++ |
| Arth-p35S-MeGI-3 | pGWB2-MeGI | — | + | + |
| Arth-p35S-MeGI-4 | pGWB2-MeGI | — | + | + |
| Arth-p35S-MeGI-5 | pGWB2-MeGI | — | — | + |
| Arth-p35S-MeGI-6 | pGWB2-MeGI | — | — | — |
| Arth-p35S-MeGI-7 | pGWB2-MeGI | — | — | + |
| Arth-p35S-MeGI-8 | pGWB2-MeGI | — | — | + |
| Arth-p35S-MeGI-9 | pGWB2-MeGI | — | — | + |
| Arth-p35S-MeGI-10 | pGWB2-MeGI | — | — | + |
| Arth-p35S-MeGI-11 | pGWB2-MeGI | + | + | ++ |
| Arth-p35S-MeGI-12 | pGWB2-MeGI | — | + | + |
| Arth-p35S-MeGI-13 | pGWB2-MeGI | — | + | + |
| Arth-p35S-MeGI-14 | pGWB2-MeGI | — | — | + |
| Arth-p35S-MeGI-15 | pGWB2-MeGI | — | — | + |
| Arth-p35S-MeGI-16 | pGWB2-MeGI | — | — | — |
| Arth-p35S-MeGI-17 | pGWB2-MeGI | — | + | NA |
| Arth-p35S-MeGI-18 | pGWB2-MeGI | + | ++ | NA |
| Arth-p35S-MeGI-19 | pGWB2-MeGI | + | + | NA |
| Arth-p35S-MeGI-20 | pGWB2-MeGI | + | + | NA |
| Arth-p35S-MeGI-21 | pGWB2-MeGI | — | — | NA |
| Arth-p35S-MeGI-22 | pGWB2-MeGI | + | ++ | NA |
| Arth-p35S-MeGI-23 | pGWB2-MeGI | + | ++ | NA |
| Arth-p35S-MeGI-24 | pGWB2-MeGI | — | — | NA |
| Arth-p35S-MeGI-25 | pGWB2-MeGI | — | — | NA |
| Arth-p35S-MeGI-26 | pGWB2-MeGI | — | — | NA |
| Arth-p35S-MeGI-27 | pGWB2-MeGI | — | + | NA |
| Arth-p35S-MeGI-28 | pGWB2-MeGI | — | + | NA |
| Arth-p35S-MeGI-29 | pGWB2-MeGI | — | + | NA |
| Arth-p35S-MeGI-30 | pGWB2-MeGI | — | — | NA |
| Arth-p35S-MeGI-31 | pGWB2-MeGI | — | + | NA |
| Arth-p35S-MeGI-32 | pGWB2-MeGI | — | + | NA |
| Arth-p35S-MeGI-33 | pGWB2-MeGI | — | — | NA |
| Arth-p35S-MeGI-34 | pGWB2-MeGI | — | — | NA |
| Arth-p35S-MeGI-35 | pGWB2-MeGI | — | + | NA |
| Arth-p35S-MeGI-36 | pGWB2-MeGI | — | — | NA |
| Arth-p35S-MeGI-37 | pGWB2-MeGI | — | — | NA |
| Arth-p35S-MeGI-38 | pGWB2-MeGI | — | — | NA |

| | | | | |
|---|---|---|---|---|
| Arth-p35S-MeGI-39 | pGWB2-MeGI | + | + | NA |
| Arth-p35S-MeGI-40 | pGWB2-MeGI | + | + | NA |
| Arth-p35S-MeGI-41 | pGWB2-MeGI | + | + | NA |
| Arth-p35S-MeGI-42 | pGWB2-MeGI | — | — | NA |
| Arth-p35S-MeGI-43 | pGWB2-MeGI | — | — | NA |
| Arth-p35S-MeGI-44 | pGWB2-MeGI | — | — | NA |
| Arth-p35S-MeGI-45 | pGWB2-MeGI | — | + | NA |
| Arth-p35S-MeGI-46 | pGWB2-MeGI | — | — | NA |
| Arth-p35S-MeGI-47 | pGWB2-MeGI | — | + | NA |
| Arth-p35S-MeGI-48 | pGWB2-MeGI | + | — | NA |
| Arth-p35S-MeGI-49 | pGWB2-MeGI | — | — | NA |
| Arth-p35S-MeGI-50 | pGWB2-MeGI | — | — | NA |
| Arth-p35S-MeGI-51 | pGWB2-MeGI | — | + | NA |
| Arth-p35S-MeGI-52 | pGWB2-MeGI | — | — | NA |
| Arth-p35S-MeGI-53 | pGWB2-MeGI | — | — | NA |
| Arth-p35S-MeGI-54 | pGWB2-MeGI | — | — | NA |
| Arth-p35S-MeGI-55 | pGWB2-MeGI | — | — | NA |
| Arth-p35S-MeGI-56 | pGWB2-MeGI | + | + | NA |
| Arth-p35S-MeGI-57 | pGWB2-MeGI | — | — | NA |
| Arth-p35S-MeGI-58 | pGWB2-MeGI | — | — | NA |
| Arth-p35S-MeGI-59 | pGWB2-MeGI | — | — | NA |
| Arth-p35S-MeGI-60 | pGWB2-MeGI | — | — | NA |
| Arth-p35S-MeGI-61 | pGWB2-MeGI | — | — | NA |
| Arth-p35S-MeGI-62 | pGWB2-MeGI | — | + | NA |
| Arth-p35S-MeGI-63 | pGWB2-MeGI | + | ++ | NA |
| Arth-p35S-MeGI-64 | pGWB2-MeGI | — | — | NA |
| Arth-p35S-MeGI-65 | pGWB2-MeGI | — | — | NA |
| Arth-p35S-MeGI-66 | pGWB2-MeGI | — | — | NA |
| Arth-p35S-MeGI-67 | pGWB2-MeGI | — | + | NA |
| Arth-p35S-MeGI-68 | pGWB2-MeGI | — | + | NA |
| Arth-p35S-MeGI-69 | pGWB2-MeGI | — | + | NA |

[a] “+” indicates feminization.

[b] “++” and “+” indicate dwarfing and semi-dwarfing, respectively (see Figure S9).

[c] Expression levels were assessed in transgenic 16 individuals by RT-PCR analysis.

### S8 Table: Phenotypic characterization of the p35S-*SiMeGI A. thaliana* transformed lines.

| T1 Line ID | introduced construct | feminization[a] | dwarfism[b] | transgene expression in whole plant[c] |
|---|---|---|---|---|
| Arth-35S-SiMeGI-1 | pGWB2-MeGI | — | — | + |
| Arth-35S-SiMeGI-2 | pGWB2-MeGI | — | — | ++ |
| Arth-35S-SiMeGI-3 | pGWB2-MeGI | — | + | ++ |
| Arth-35S-SiMeGI-4 | pGWB2-MeGI | — | — | ++ |
| Arth-35S-SiMeGI-5 | pGWB2-MeGI | — | + | + |
| Arth-35S-SiMeGI-6 | pGWB2-MeGI | — | — | + |
| Arth-35S-SiMeGI-7 | pGWB2-MeGI | — | + | ++ |
| Arth-35S-SiMeGI-8 | pGWB2-MeGI | — | — | ++ |
| Arth-35S-SiMeGI-9 | pGWB2-MeGI | — | — | + |
| Arth-35S-SiMeGI-10 | pGWB2-MeGI | — | — | + |
| Arth-35S-SiMeGI-11 | pGWB2-MeGI | — | — | ++ |
| Arth-35S-SiMeGI-12 | pGWB2-MeGI | + | + | ++ |
| Arth-35S-SiMeGI-13 | pGWB2-MeGI | — | — | + |
| Arth-35S-SiMeGI-14 | pGWB2-MeGI | — | — | + |
| Arth-35S-SiMeGI-15 | pGWB2-MeGI | — | — | + |
| Arth-35S-SiMeGI-16 | pGWB2-MeGI | — | — | + |
| Arth-35S-SiMeGI-17 | pGWB2-MeGI | — | — | + |
| Arth-35S-SiMeGI-18 | pGWB2-MeGI | + | + | + |
| Arth-35S-SiMeGI-19 | pGWB2-MeGI | — | + | + |
| Arth-35S-SiMeGI-20 | pGWB2-MeGI | — | — | + |
| Arth-35S-SiMeGI-21 | pGWB2-MeGI | — | — | + |
| Arth-35S-SiMeGI-22 | pGWB2-MeGI | — | + | + |
| Arth-35S-SiMeGI-23 | pGWB2-MeGI | — | — | + |
| Arth-35S-SiMeGI-24 | pGWB2-MeGI | — | — | + |
| Arth-35S-SiMeGI-25 | pGWB2-MeGI | — | + | NA |
| Arth-35S-SiMeGI-26 | pGWB2-MeGI | — | + | NA |
| Arth-35S-SiMeGI-27 | pGWB2-MeGI | — | — | NA |
| Arth-35S-SiMeGI-28 | pGWB2-MeGI | — | — | NA |
| Arth-35S-SiMeGI-29 | pGWB2-MeGI | — | — | NA |
| Arth-35S-SiMeGI-30 | pGWB2-MeGI | — | — | NA |
| Arth-35S-SiMeGI-31 | pGWB2-MeGI | — | — | NA |
| Arth-35S-SiMeGI-32 | pGWB2-MeGI | — | — | NA |
| Arth-35S-SiMeGI-33 | pGWB2-MeGI | — | — | NA |
| Arth-35S-SiMeGI-34 | pGWB2-MeGI | — | — | NA |
| Arth-35S-SiMeGI-35 | pGWB2-MeGI | — | — | NA |
| Arth-35S-SiMeGI-36 | pGWB2-MeGI | — | — | NA |
| Arth-35S-SiMeGI-37 | pGWB2-MeGI | — | + | NA |
| Arth-35S-SiMeGI-38 | pGWB2-MeGI | — | — | NA |
| Arth-35S-SiMeGI-39 | pGWB2-MeGI | — | — | NA |

| | | | | |
|---|---|---|---|---|
| Arth-35S-SiMeGI-40 | pGWB2-MeGI | — | — | NA |
| Arth-35S-SiMeGI-41 | pGWB2-MeGI | — | — | NA |
| Arth-35S-SiMeGI-42 | pGWB2-MeGI | — | — | NA |
| Arth-35S-SiMeGI-43 | pGWB2-MeGI | — | + | NA |
| Arth-35S-SiMeGI-44 | pGWB2-MeGI | — | + | NA |
| Arth-35S-SiMeGI-45 | pGWB2-MeGI | — | — | NA |
| Arth-35S-SiMeGI-46 | pGWB2-MeGI | — | + | NA |
| Arth-35S-SiMeGI-47 | pGWB2-MeGI | — | + | NA |
| Arth-35S-SiMeGI-48 | pGWB2-MeGI | — | — | NA |
| Arth-35S-SiMeGI-49 | pGWB2-MeGI | — | — | NA |
| Arth-35S-SiMeGI-50 | pGWB2-MeGI | — | — | NA |
| Arth-35S-SiMeGI-51 | pGWB2-MeGI | — | — | NA |
| Arth-35S-SiMeGI-52 | pGWB2-MeGI | — | + | NA |
| Arth-35S-SiMeGI-53 | pGWB2-MeGI | — | — | NA |

[a] "+" indicates feminization.
[b] "++" and "+" indicate dwarfing and semi-dwarfing, respectively (see Figure S9).
[c] Expression levels were assessed in 16 transgenic individuals by RT-PCR analysis.

### S9 Table: Plant materials

| name | Gender[a] | Generation/population | Assessment[b] | Cross (year) |
|---|---|---|---|---|
| Kunsenshi-male | Male | Male parent of the KK amd VM populations | RAD/GBS, draft genome sequencing, transcriptome | |
| Kunsenshi-female | Female | Female parent of the KK population | RAD/GBS, transcriptome | |
| Budogaki | Female | Female parent of the VM population | RAD/na | |
| KK1_001 | Male | F1, KK population | RAD/GBS | 2004 |
| KK1_002 | Male | F1, KK population | RAD/GBS | 2004 |
| KK1_003 | Male | F1, KK population | RAD/GBS | 2004 |
| KK1_004 | Male | F1, KK population | RAD/GBS | 2004 |
| KK1_005 | Male | F1, KK population | RAD/na | 2004 |
| KK1_006 | Male | F1, KK population | RAD/GBS | 2004 |
| KK1_007 | Female | F1, KK population | na/GBS | 2004 |
| KK1_009 | Male | F1, KK population | RAD/GBS | 2004 |
| KK1_010 | Female | F1, KK population | RAD/GBS | 2004 |
| KK1_011 | Female | F1, KK population | RAD/GBS | 2004 |
| KK1_012 | Male | F1, KK population | RAD/GBS | 2004 |
| KK1_013 | Female | F1, KK population | RAD/na | 2004 |
| KK1_014 | Female | F1, KK population | RAD/GBS | 2004 |
| KK1_016 | Female | F1, KK population | RAD/GBS | 2004 |
| KK1_017 | Female | F1, KK population | RAD/GBS | 2004 |
| KK1_018 | Female | F1, KK population | RAD/na | 2004 |
| KK1_020 | Female | F1, KK population | RAD/GBS | 2004 |
| KK1_021 | Male | F1, KK population | RAD/na | 2004 |
| KK1_022 | Male | F1, KK population | RAD/GBS | 2004 |
| KK1_023 | Female | F1, KK population | RAD/GBS | 2004 |
| KK1_024 | Female | F1, KK population | RAD/GBS | 2004 |
| KK1_025 | Male | F1, KK population | RAD/GBS | 2004 |
| KK1_027 | Female | F1, KK population | RAD/GBS | 2004 |
| KK1_028 | Male | F1, KK population | RAD/GBS | 2004 |
| KK1_029 | Female | F1, KK population | RAD/GBS | 2004 |
| KK1_030 | Female | F1, KK population | na/GBS | 2004 |
| KK1_031 | Male | F1, KK population | RAD/GBS | 2004 |
| KK1_032 | Male | F1, KK population | RAD/GBS | 2004 |
| KK1_033 | Female | F1, KK population | RAD/GBS | 2004 |
| KK1_034 | Male | F1, KK population | RAD/GBS | 2004 |
| KK1_035 | Female | F1, KK population | RAD/GBS | 2004 |
| KK1_036 | Male | F1, KK population | RAD/GBS | 2004 |
| KK1_037 | Female | F1, KK population | na/GBS | 2004 |
| KK1_038 | Female | F1, KK population | RAD/GBS | 2004 |
| KK1_039 | Female | F1, KK population | RAD/GBS | 2004 |
| KK1_040 | Male | F1, KK population | RAD/GBS | 2004 |
| KK1_041 | Female | F1, KK population | RAD/GBS | 2004 |
| KK1_042 | Female | F1, KK population | RAD/na | 2004 |
| KK1_043 | Male | F1, KK population | RAD/na | 2004 |
| KK1_044 | Female | F1, KK population | RAD/na | 2004 |
| KK1_045 | Female | F1, KK population | RAD/na | 2004 |
| KK1_046 | Female | F1, KK population | RAD/GBS | 2004 |
| KK1_047 | Female | F1, KK population | RAD/na | 2004 |
| KK1_048 | Male | F1, KK population | RAD/GBS | 2004 |
| KK1_049 | Female | F1, KK population | RAD/GBS | 2004 |
| KK1_051 | Female | F1, KK population | RAD/GBS | 2004 |
| KK1_052 | Male | F1, KK population | RAD/GBS | 2004 |
| KK1_054 | Female | F1, KK population | RAD/GBS | 2004 |
| KK1_055 | Female | F1, KK population | RAD/na | 2004 |
| KK1_056 | Female | F1, KK population | RAD/GBS | 2004 |

| | | | | |
|---|---|---|---|---|
| KK1_057 | Male | F1, KK population | RAD/na | 2004 |
| KK1_058 | Female | F1, KK population | RAD/na | 2004 |
| KK1_059 | Female | F1, KK population | RAD/GBS | 2004 |
| KK1_060 | Female | F1, KK population | RAD/GBS | 2004 |
| KK1_061 | Male | F1, KK population | RAD/GBS | 2004 |
| KK1_062 | Female | F1, KK population | RAD/GBS | 2004 |
| KK1_063 | Male | F1, KK population | RAD/GBS | 2004 |
| KK1_064 | Female | F1, KK population | RAD/GBS | 2004 |
| KK2_001 | Female | F1, KK population | RAD/na | 2011 |
| KK2_002 | Female | F1, KK population | RAD/GBS | 2011 |
| KK2_003 | Male | F1, KK population | RAD/na | 2011 |
| KK2_004 | NF | F1, KK population | RAD/GBS | 2011 |
| KK2_005 | Female | F1, KK population | RAD/GBS | 2011 |
| KK2_006 | NF | F1, KK population | RAD/GBS | 2011 |
| KK2_007 | Male | F1, KK population | RAD/na | 2011 |
| KK2_008 | Male | F1, KK population | RAD/GBS | 2011 |
| KK2_009 | Male | F1, KK population | RAD/GBS | 2011 |
| KK2_010 | Female | F1, KK population | RAD/GBS | 2011 |
| KK2_011 | Female | F1, KK population | RAD/GBS | 2011 |
| KK2_012 | NF | F1, KK population | RAD/GBS | 2011 |
| KK2_013 | Male | F1, KK population | RAD/GBS | 2011 |
| KK2_014 | NF | F1, KK population | RAD/GBS | 2011 |
| KK2_015 | Male | F1, KK population | RAD/GBS | 2011 |
| KK2_016 | Male | F1, KK population | RAD/GBS | 2011 |
| KK2_017 | NF | F1, KK population | RAD/GBS | 2011 |
| KK2_018 | Female | F1, KK population | RAD/GBS | 2011 |
| KK2_019 | Female | F1, KK population | RAD/GBS | 2011 |
| KK2_020 | Male | F1, KK population | RAD/GBS | 2011 |
| KK2_021 | NF | F1, KK population | RAD/GBS | 2011 |
| KK2_022 | NF | F1, KK population | RAD/GBS | 2011 |
| KK2_023 | Female | F1, KK population | RAD/na | 2011 |
| KK2_024 | Female | F1, KK population | RAD/GBS | 2011 |
| KK2_025 | Male | F1, KK population | RAD/GBS | 2011 |
| KK2_026 | Male | F1, KK population | RAD/GBS | 2011 |
| KK2_027 | Female | F1, KK population | RAD/GBS | 2011 |
| KK2_028 | Female | F1, KK population | RAD/GBS | 2011 |
| KK2_029 | Female | F1, KK population | RAD/GBS | 2011 |
| KK2_030 | Female | F1, KK population | RAD/GBS | 2011 |
| KK2_031 | Female | F1, KK population | RAD/GBS | 2011 |
| KK2_032 | Female | F1, KK population | RAD/GBS | 2011 |
| KK2_033 | Female | F1, KK population | RAD/GBS | 2011 |
| KK2_034 | Female | F1, KK population | RAD/GBS | 2011 |
| KK2_035 | Male | F1, KK population | RAD/GBS | 2011 |
| KK2_036 | Female | F1, KK population | RAD/na | 2011 |
| KK2_037 | Male | F1, KK population | RAD/GBS | 2011 |
| KK2_038 | Female | F1, KK population | RAD/GBS | 2011 |
| KK2_039 | Male | F1, KK population | RAD/GBS | 2011 |
| KK2_040 | Female | F1, KK population | RAD/GBS | 2011 |
| KK2_041 | Male | F1, KK population | RAD/GBS | 2011 |
| KK2_042 | NF | F1, KK population | RAD/GBS | 2011 |
| KK2_043 | Male | F1, KK population | RAD/GBS | 2011 |
| KK2_044 | Female | F1, KK population | RAD/GBS | 2011 |
| KK2_045 | NF | F1, KK population | RAD/GBS | 2011 |
| KK2_046 | Female | F1, KK population | RAD/GBS | 2011 |
| KK2_047 | Male | F1, KK population | RAD/GBS | 2011 |
| KK2_048 | Female | F1, KK population | RAD/GBS | 2011 |
| KK2_049 | Female | F1, KK population | RAD/GBS | 2011 |
| KK2_050 | Female | F1, KK population | RAD/GBS | 2011 |
| KK2_051 | Male | F1, KK population | RAD/GBS | 2011 |
| KK2_052 | Female | F1, KK population | RAD/GBS | 2011 |
| KK2_053 | Male | F1, KK population | RAD/GBS | 2011 |
| KK2_054 | Female | F1, KK population | RAD/GBS | 2011 |
| KK2_055 | Female | F1, KK population | RAD/GBS | 2011 |
| KK2_056 | Female | F1, KK population | RAD/GBS | 2011 |
| KK2_057 | Male | F1, KK population | RAD/GBS | 2011 |

| | | | | |
|---|---|---|---|---|
| KK2_058 | Male | F1, KK population | RAD/GBS | 2011 |
| KK2_059 | NF | F1, KK population | RAD/GBS | 2011 |
| KK2_060 | Male | F1, KK population | RAD/GBS | 2011 |
| KK2_061 | Female | F1, KK population | RAD/GBS | 2011 |
| KK2_062 | NF | F1, KK population | RAD/GBS | 2011 |
| KK2_063 | Female | F1, KK population | RAD/GBS | 2011 |
| KK2_064 | NF | F1, KK population | RAD/GBS | 2011 |
| KK2_065 | NF | F1, KK population | RAD/GBS | 2011 |
| KK2_066 | NF | F1, KK population | RAD/na | 2011 |
| KK2_067 | NF | F1, KK population | RAD/GBS | 2011 |
| KK2_068 | NF | F1, KK population | RAD/na | 2011 |
| KK2_069 | NF | F1, KK population | RAD/na | 2011 |
| KK2_070 | NF | F1, KK population | RAD/na | 2011 |
| KK2_071 | Male | F1, KK population | RAD/GBS | 2011 |
| KK2_072 | Male | F1, KK population | RAD/na | 2011 |
| KK2_073 | Male | F1, KK population | RAD/na | 2011 |
| KK2_074 | Male | F1, KK population | RAD/GBS | 2011 |
| KK2_075 | Female | F1, KK population | RAD/na | 2011 |
| KK2_076 | Male | F1, KK population | RAD/GBS | 2011 |
| KK2_077 | NF | F1, KK population | RAD/GBS | 2011 |
| KK2_078 | Female | F1, KK population | RAD/GBS | 2011 |
| KK2_079 | Female | F1, KK population | RAD/GBS | 2011 |
| KK2_080 | NF | F1, KK population | RAD/GBS | 2011 |
| KK2_081 | NF | F1, KK population | RAD/GBS | 2011 |
| KK2_085 | Male | F1, KK population | na/GBS | 2011 |
| KK2_086 | Male | F1, KK population | na/GBS | 2011 |
| KK2_088 | Male | F1, KK population | na/GBS | 2011 |
| KK2_089 | Female | F1, KK population | na/GBS | 2011 |
| KK2_090 | Male | F1, KK population | na/GBS | 2011 |
| KK2_091 | NF | F1, KK population | na/GBS | 2011 |
| KK2_101 | Male | F1, KK population | RAD/na | 2011 |
| KK2_104 | Male | F1, KK population | RAD/GBS | 2011 |
| KK2_105 | Male | F1, KK population | RAD/GBS | 2011 |
| KK2_106 | NF | F1, KK population | RAD/GBS | 2011 |
| KK2_107 | Female | F1, KK population | RAD/na | 2011 |
| KK2_108 | NF | F1, KK population | RAD/GBS | 2011 |
| KK2_109 | NF | F1, KK population | RAD/na | 2011 |
| KK2_111 | NF | F1, KK population | na/GBS | 2011 |
| KK2_112 | NF | F1, KK population | RAD/na | 2011 |
| KK2_113 | Male | F1, KK population | RAD/GBS | 2011 |
| KK2_114 | NF | F1, KK population | RAD/na | 2011 |
| KK2_117 | NF | F1, KK population | RAD/GBS | 2011 |
| KK2_118 | NF | F1, KK population | RAD/GBS | 2011 |
| KK2_119 | NF | F1, KK population | RAD/GBS | 2011 |
| KK2_120 | Female | F1, KK population | RAD/GBS | 2011 |
| KK2_121 | Male | F1, KK population | RAD/na | 2011 |
| KK2_122 | NF | F1, KK population | RAD/GBS | 2011 |
| KK2_123 | NF | F1, KK population | na/GBS | 2011 |
| KK2_124 | NF | F1, KK population | RAD/GBS | 2011 |
| KK2_125 | NF | F1, KK population | RAD/GBS | 2011 |
| KK2_126 | NF | F1, KK population | RAD/GBS | 2011 |
| KK2_128 | NF | F1, KK population | RAD/GBS | 2011 |
| KK2_129 | NF | F1, KK population | RAD/na | 2011 |
| KK2_130 | NF | F1, KK population | RAD/GBS | 2011 |
| KK2_131 | Male | F1, KK population | RAD/GBS | 2011 |
| KK2_132 | NF | F1, KK population | RAD/na | 2011 |
| KK2_133 | NF | F1, KK population | RAD/GBS | 2011 |
| KK2_134 | NF | F1, KK population | RAD/GBS | 2011 |
| KK2_135 | NF | F1, KK population | RAD/GBS | 2011 |
| KK2_136 | NF | F1, KK population | RAD/GBS | 2011 |
| KK2_137 | NF | F1, KK population | RAD/GBS | 2011 |
| KK2_138 | NF | F1, KK population | na/GBS | 2011 |
| KK2_139 | NF | F1, KK population | na/GBS | 2011 |
| KK2_140 | NF | F1, KK population | RAD/GBS | 2011 |
| KK2_141 | NF | F1, KK population | RAD/GBS | 2011 |

| | | | | |
|---|---|---|---|---|
| KK2_142 | NF | F1, KK population | RAD/GBS | 2011 |
| KK2_143 | NF | F1, KK population | RAD/GBS | 2011 |
| KK2_144 | Female | F1, KK population | RAD/GBS | 2011 |
| KK2_145 | NF | F1, KK population | RAD/GBS | 2011 |
| KK2_146 | NF | F1, KK population | RAD/GBS | 2011 |
| KK2_147 | NF | F1, KK population | RAD/GBS | 2011 |
| KK2_148 | NF | F1, KK population | RAD/GBS | 2011 |
| KK2_149 | Male | F1, KK population | RAD/GBS | 2011 |
| KK2_150 | NF | F1, KK population | RAD/GBS | 2011 |
| KK2_151 | NF | F1, KK population | RAD/GBS | 2011 |
| KK2_152 | Male | F1, KK population | RAD/GBS | 2011 |
| KK2_154 | NF | F1, KK population | RAD/GBS | 2011 |
| KK2_155 | NF | F1, KK population | RAD/GBS | 2011 |
| KK2_156 | NF | F1, KK population | RAD/GBS | 2011 |
| KK2_157 | Female | F1, KK population | RAD/GBS | 2011 |
| KK2_158 | Male | F1, KK population | RAD/GBS | 2011 |
| KK2_159 | Male | F1, KK population | RAD/GBS | 2011 |
| KK2_160 | Male | F1, KK population | RAD/GBS | 2011 |
| KK2_161 | NF | F1, KK population | RAD/GBS | 2011 |
| KK2_162 | Male | F1, KK population | RAD/GBS | 2011 |
| KK2_163 | NF | F1, KK population | RAD/GBS | 2011 |
| KK2_164 | NF | F1, KK population | RAD/GBS | 2011 |
| KK2_165 | Female | F1, KK population | RAD/GBS | 2011 |
| KK2_166 | Female | F1, KK population | RAD/na | 2011 |
| KK2_167 | NF | F1, KK population | RAD/GBS | 2011 |
| KK2_168 | NF | F1, KK population | RAD/GBS | 2011 |
| KK2_169 | Female | F1, KK population | RAD/GBS | 2011 |
| KK2_170 | NF | F1, KK population | RAD/GBS | 2011 |
| KK2_171 | Male | F1, KK population | RAD/GBS | 2011 |
| KK2_172 | NF | F1, KK population | RAD/GBS | 2011 |
| KK2_173 | Female | F1, KK population | RAD/GBS | 2011 |
| KK2_174 | NF | F1, KK population | RAD/GBS | 2011 |
| KK2_175 | Male | F1, KK population | RAD/GBS | 2011 |
| KK2_176 | Male | F1, KK population | RAD/GBS | 2011 |
| KK2_177 | NF | F1, KK population | RAD/na | 2011 |
| KK2_178 | NF | F1, KK population | RAD/GBS | 2011 |
| KK2_179 | Male | F1, KK population | RAD/na | 2011 |
| KK2_180 | NF | F1, KK population | RAD/na | 2011 |
| KK2_181 | Male | F1, KK population | RAD/GBS | 2011 |
| KK2_182 | NF | F1, KK population | RAD/GBS | 2011 |
| KK2_183 | NF | F1, KK population | RAD/GBS | 2011 |
| KK2_184 | NF | F1, KK population | RAD/GBS | 2011 |
| KK2_185 | Female | F1, KK population | RAD/GBS | 2011 |
| KK2_186 | Male | F1, KK population | RAD/GBS | 2011 |
| KK2_187 | NF | F1, KK population | RAD/GBS | 2011 |
| KK2_188 | NF | F1, KK population | RAD/GBS | 2011 |
| KK2_189 | NF | F1, KK population | RAD/GBS | 2011 |
| KK2_190 | Male | F1, KK population | RAD/GBS | 2011 |
| KK2_191 | NF | F1, KK population | RAD/GBS | 2011 |
| KK2_192 | NF | F1, KK population | RAD/GBS | 2011 |
| KK2_193 | NF | F1, KK population | RAD/GBS | 2011 |
| KK2_194 | NF | F1, KK population | RAD/GBS | 2011 |
| KK2_195 | Female | F1, KK population | RAD/GBS | 2011 |
| KK2_196 | NF | F1, KK population | RAD/GBS | 2011 |
| KK2_197 | Female | F1, KK population | RAD/GBS | 2011 |
| KK2_198 | NF | F1, KK population | RAD/GBS | 2011 |
| KK2_199 | Female | F1, KK population | RAD/GBS | 2011 |
| KK2_200 | Male | F1, KK population | RAD/GBS | 2011 |
| KK2_201 | Female | F1, KK population | RAD/GBS | 2011 |
| KK2_202 | NF | F1, KK population | RAD/GBS | 2011 |
| KK2_203 | NF | F1, KK population | RAD/GBS | 2011 |
| KK2_204 | Female | F1, KK population | RAD/GBS | 2011 |
| KK2_205 | Male | F1, KK population | RAD/GBS | 2011 |
| KK2_206 | Male | F1, KK population | RAD/GBS | 2011 |
| KK2_207 | Male | F1, KK population | RAD/GBS | 2011 |

| | | | | |
|---|---|---|---|---|
| KK2_208 | NF | F1, KK population | RAD/GBS | 2011 |
| KK2_209 | NF | F1, KK population | RAD/GBS | 2011 |
| KK2_210 | Female | F1, KK population | RAD/GBS | 2011 |
| KK2_211 | NF | F1, KK population | RAD/GBS | 2011 |
| KK2_212 | NF | F1, KK population | RAD/GBS | 2011 |
| KK2_213 | Female | F1, KK population | RAD/GBS | 2011 |
| KK2_214 | Female | F1, KK population | RAD/GBS | 2011 |
| KK2_215 | NF | F1, KK population | RAD/GBS | 2011 |
| KK2_216 | Male | F1, KK population | RAD/GBS | 2011 |
| KK2_217 | Male | F1, KK population | RAD/GBS | 2011 |
| KK2_218 | Male | F1, KK population | RAD/GBS | 2011 |
| KK2_219 | Male | F1, KK population | RAD/GBS | 2011 |
| KK2_220 | Male | F1, KK population | RAD/GBS | 2011 |
| KK2_221 | Male | F1, KK population | RAD/GBS | 2011 |
| KK2_222 | Female | F1, KK population | RAD/na | 2011 |
| KK2_223 | NF | F1, KK population | RAD/GBS | 2011 |
| KK2_224 | Female | F1, KK population | RAD/GBS | 2011 |
| KK2_225 | Male | F1, KK population | RAD/na | 2011 |
| KK2_226 | Male | F1, KK population | RAD/GBS | 2011 |
| KK2_227 | NF | F1, KK population | RAD/GBS | 2011 |
| KK2_228 | NF | F1, KK population | RAD/GBS | 2011 |
| KK2_229 | Female | F1, KK population | RAD/GBS | 2011 |
| KK2_230 | NF | F1, KK population | RAD/GBS | 2011 |
| KK2_231 | NF | F1, KK population | RAD/na | 2011 |
| KK2_232 | NF | F1, KK population | RAD/GBS | 2011 |
| KK2_233 | Female | F1, KK population | RAD/GBS | 2011 |
| KK2_234 | Female | F1, KK population | RAD/GBS | 2011 |
| KK2_235 | NF | F1, KK population | RAD/GBS | 2011 |
| KK2_236 | NF | F1, KK population | RAD/GBS | 2011 |
| KK2_237 | NF | F1, KK population | RAD/GBS | 2011 |
| KK2_238 | NF | F1, KK population | RAD/GBS | 2011 |
| KK2_239 | NF | F1, KK population | RAD/GBS | 2011 |
| KK2_240 | Female | F1, KK population | RAD/GBS | 2011 |
| KK2_241 | Male | F1, KK population | RAD/GBS | 2011 |
| KK2_242 | Male | F1, KK population | RAD/na | 2011 |
| KK2_243 | Male | F1, KK population | RAD/na | 2011 |
| KK2_244 | NF | F1, KK population | RAD/GBS | 2011 |
| KK2_245 | NF | F1, KK population | RAD/na | 2011 |
| KK2_246 | NF | F1, KK population | RAD/GBS | 2011 |
| KK2_247 | Male | F1, KK population | RAD/GBS | 2011 |
| KK2_248 | NF | F1, KK population | RAD/GBS | 2011 |
| KK2_249 | Male | F1, KK population | RAD/GBS | 2011 |
| KK2_250 | NF | F1, KK population | RAD/GBS | 2011 |
| KK2_251 | Male | F1, KK population | RAD/na | 2011 |
| KK2_252 | Female | F1, KK population | RAD/GBS | 2011 |
| KK2_253 | Female | F1, KK population | RAD/GBS | 2011 |
| KK2_254 | Female | F1, KK population | RAD/GBS | 2011 |
| KK2_255 | NF | F1, KK population | RAD/na | 2011 |
| KK2_256 | NF | F1, KK population | RAD/GBS | 2011 |
| KK2_257 | NF | F1, KK population | RAD/GBS | 2011 |
| KK2_258 | NF | F1, KK population | RAD/GBS | 2011 |
| KK2_259 | NF | F1, KK population | RAD/GBS | 2011 |
| KK2_260 | Male | F1, KK population | RAD/GBS | 2011 |
| KK2_261 | Female | F1, KK population | RAD/GBS | 2011 |
| KK2_262 | NF | F1, KK population | na/GBS | 2011 |
| KK2_264 | NF | F1, KK population | na/GBS | 2011 |
| KK2_265 | NF | F1, KK population | na/GBS | 2011 |
| KK2_266 | Male | F1, KK population | na/GBS | 2011 |
| KK2_267 | NF | F1, KK population | na/GBS | 2011 |
| KK2_268 | NF | F1, KK population | na/GBS | 2011 |
| KK2_269 | Male | F1, KK population | na/GBS | 2011 |
| KK2_270 | Female | F1, KK population | na/GBS | 2011 |
| KK2_271 | NF | F1, KK population | na/GBS | 2011 |
| KK2_272 | NF | F1, KK population | na/GBS | 2011 |
| KK2_273 | NF | F1, KK population | na/GBS | 2011 |

| | | | | |
|---|---|---|---|---|
| KK2_274 | NF | F1, KK population | na/GBS | 2011 |
| KK2_275 | NF | F1, KK population | na/GBS | 2011 |
| KK2_276 | NF | F1, KK population | na/GBS | 2011 |
| KK2_277 | NF | F1, KK population | na/GBS | 2011 |
| VM_1 | NA | F1, VM population | RAD/na | 2009 |
| VM_2 | NA | F1, VM population | RAD/na | 2009 |
| VM_3 | NA | F1, VM population | RAD/na | 2009 |
| VM_4 | NA | F1, VM population | RAD/na | 2009 |
| VM_5 | NA | F1, VM population | RAD/na | 2009 |
| VM_6 | NA | F1, VM population | RAD/na | 2009 |
| VM_7 | NA | F1, VM population | RAD/na | 2009 |
| VM_8 | NA | F1, VM population | RAD/na | 2009 |
| VM_9 | NA | F1, VM population | RAD/na | 2009 |
| VM_10 | NA | F1, VM population | RAD/na | 2009 |
| VM_11 | NA | F1, VM population | RAD/na | 2009 |
| VM_12 | NA | F1, VM population | RAD/na | 2009 |
| VM_13 | NA | F1, VM population | RAD/na | 2009 |
| VM_14 | NA | F1, VM population | RAD/na | 2009 |
| VM_15 | NA | F1, VM population | RAD/na | 2009 |
| VM_16 | NA | F1, VM population | RAD/na | 2009 |
| VM_17 | NA | F1, VM population | RAD/na | 2009 |
| VM_18 | NA | F1, VM population | RAD/na | 2009 |
| VM_19 | NA | F1, VM population | RAD/na | 2009 |
| VM_20 | NA | F1, VM population | RAD/na | 2009 |
| VM_21 | NA | F1, VM population | RAD/na | 2009 |
| VM_22 | NA | F1, VM population | RAD/na | 2009 |
| VM_23 | NA | F1, VM population | RAD/na | 2009 |
| VM_24 | NA | F1, VM population | RAD/na | 2009 |
| VM_25 | NA | F1, VM population | RAD/na | 2009 |
| VM_26 | NA | F1, VM population | RAD/na | 2009 |
| VM_27 | NA | F1, VM population | RAD/na | 2009 |
| VM_28 | NA | F1, VM population | RAD/na | 2009 |
| VM_29 | NA | F1, VM population | RAD/na | 2009 |
| VM_30 | NA | F1, VM population | RAD/na | 2009 |
| VM_31 | NA | F1, VM population | RAD/na | 2009 |
| VM_32 | NA | F1, VM population | RAD/na | 2009 |
| VM_33 | NA | F1, VM population | RAD/na | 2009 |
| VM_34 | NA | F1, VM population | RAD/na | 2009 |
| VM_35 | NA | F1, VM population | RAD/na | 2009 |
| VM_36 | NA | F1, VM population | RAD/na | 2009 |
| VM_37 | NA | F1, VM population | RAD/na | 2009 |
| VM_38 | NA | F1, VM population | RAD/na | 2009 |
| VM_39 | NA | F1, VM population | RAD/na | 2009 |
| VM_40 | NA | F1, VM population | RAD/na | 2009 |
| VM_41 | NA | F1, VM population | RAD/na | 2009 |
| VM_42 | NA | F1, VM population | RAD/na | 2009 |
| VM_43 | NA | F1, VM population | RAD/na | 2009 |
| VM_44 | NA | F1, VM population | RAD/na | 2009 |
| VM_45 | NA | F1, VM population | RAD/na | 2009 |
| VM_46 | NA | F1, VM population | RAD/na | 2009 |
| VM_47 | NA | F1, VM population | RAD/na | 2009 |
| VM_48 | NA | F1, VM population | RAD/na | 2009 |
| VM_49 | NA | F1, VM population | RAD/na | 2009 |
| VM_50 | NA | F1, VM population | RAD/na | 2009 |
| VM_51 | NA | F1, VM population | RAD/na | 2009 |
| VM_52 | NA | F1, VM population | RAD/na | 2009 |
| VM_53 | NA | F1, VM population | RAD/na | 2009 |
| VM_54 | NA | F1, VM population | RAD/na | 2009 |
| VM_55 | NA | F1, VM population | RAD/na | 2009 |
| VM_56 | NA | F1, VM population | RAD/na | 2009 |
| VM_57 | NA | F1, VM population | RAD/na | 2009 |
| VM_58 | NA | F1, VM population | RAD/na | 2009 |
| VM_59 | NA | F1, VM population | RAD/na | 2009 |
| VM_60 | NA | F1, VM population | RAD/na | 2009 |
| VM_61 | NA | F1, VM population | RAD/na | 2009 |

| | | | | |
|---|---|---|---|---|
| VM_62 | NA | F1, VM population | RAD/na | 2009 |
| VM_63 | NA | F1, VM population | RAD/na | 2009 |
| VM_64 | NA | F1, VM population | RAD/na | 2009 |
| VM_65 | NA | F1, VM population | RAD/na | 2009 |
| VM_66 | NA | F1, VM population | RAD/na | 2009 |
| VM_67 | NA | F1, VM population | RAD/na | 2009 |
| VM_68 | NA | F1, VM population | RAD/na | 2009 |
| VM_69 | NA | F1, VM population | RAD/na | 2009 |
| VM_70 | NA | F1, VM population | RAD/na | 2009 |
| VM_71 | NA | F1, VM population | RAD/na | 2009 |
| VM_72 | NA | F1, VM population | RAD/na | 2009 |
| VM_73 | NA | F1, VM population | RAD/na | 2009 |
| VM_74 | NA | F1, VM population | RAD/na | 2009 |
| VM_75 | NA | F1, VM population | RAD/na | 2009 |
| VM_76 | NA | F1, VM population | RAD/na | 2009 |
| VM_77 | NA | F1, VM population | RAD/na | 2009 |
| VM_78 | NA | F1, VM population | RAD/na | 2009 |
| VM_79 | NA | F1, VM population | RAD/na | 2009 |
| VM_80 | NA | F1, VM population | RAD/na | 2009 |
| VM_81 | NA | F1, VM population | RAD/na | 2009 |
| VM_82 | NA | F1, VM population | RAD/na | 2009 |
| VM_83 | NA | F1, VM population | RAD/na | 2009 |
| VM_84 | NA | F1, VM population | RAD/na | 2009 |
| VM_85 | NA | F1, VM population | RAD/na | 2009 |
| VM_86 | NA | F1, VM population | RAD/na | 2009 |
| VM_87 | NA | F1, VM population | RAD/na | 2009 |
| VM_88 | NA | F1, VM population | RAD/na | 2009 |
| VM_89 | NA | F1, VM population | RAD/na | 2009 |
| VM_90 | NA | F1, VM population | RAD/na | 2009 |
| VM_91 | NA | F1, VM population | RAD/na | 2009 |
| VM_92 | NA | F1, VM population | RAD/na | 2009 |
| VM_93 | NA | F1, VM population | RAD/na | 2009 |
| VM_94 | NA | F1, VM population | RAD/na | 2009 |
| VM_95 | NA | F1, VM population | RAD/na | 2009 |
| VM_96 | NA | F1, VM population | RAD/na | 2009 |
| VM_97 | NA | F1, VM population | RAD/na | 2009 |
| VM_98 | NA | F1, VM population | RAD/na | 2009 |
| VM_99 | NA | F1, VM population | RAD/na | 2009 |
| VM_100 | NA | F1, VM population | RAD/na | 2009 |
| VM_101 | NA | F1, VM population | RAD/na | 2009 |
| VM_102 | NA | F1, VM population | RAD/na | 2009 |
| VM_103 | NA | F1, VM population | RAD/na | 2009 |
| VM_104 | NA | F1, VM population | RAD/na | 2009 |
| VM_105 | NA | F1, VM population | RAD/na | 2009 |
| VM_106 | NA | F1, VM population | RAD/na | 2009 |
| VM_107 | NA | F1, VM population | RAD/na | 2009 |
| VM_108 | NA | F1, VM population | RAD/na | 2009 |
| VM_109 | NA | F1, VM population | RAD/na | 2009 |
| VM_110 | NA | F1, VM population | RAD/na | 2009 |
| VM_111 | NA | F1, VM population | RAD/na | 2009 |
| VM_112 | NA | F1, VM population | RAD/na | 2009 |
| VM_113 | NA | F1, VM population | RAD/na | 2009 |
| VM_114 | NA | F1, VM population | RAD/na | 2009 |
| VM_115 | NA | F1, VM population | RAD/na | 2009 |
| VM_116 | NA | F1, VM population | RAD/na | 2009 |
| VM_117 | NA | F1, VM population | RAD/na | 2009 |
| VM_118 | NA | F1, VM population | RAD/na | 2009 |
| VM_119 | NA | F1, VM population | RAD/na | 2009 |

[a] NF: not yet flowered in 2017, NA: not analyzed.

[b] Information of both RAD (ddRAD-Seq analysis) and GBS were given. na: not analyzed.

### References

M. D. Bennett, P. Bhandol, I. J. Leitch, Nuclear DNA amounts in angiosperms and their modern uses - 807 new estimates. *Ann. Bot.* **86**, 859-909 (2000).

C. Y. Cheng, *et al*. Araport11: a complete reannotation of the Arabidopsis thaliana reference genome. *Plant J.* **89**, 789-804 (2017).

L. Li, C. J. Stoeckert, D. S. Roos, OrthoMCL: identification of ortholog groups for eukaryotic genomes. *Genome Res.* **13**, 2178-2189 (2003).

I. Verde, *et al*. The high-quality draft genome of peach (*Prunus persica*) identifies unique patterns of genetic diversity, domestication and genome evolution. *Nat. Genet.* **45**, 487-494 (2013).

R. Ming, *et al*. The draft genome of the transgenic tropical fruit tree papaya (*Carica papaya* Linnaeus). *Nature* **452**, 991-996 (2008).
